# The AusTraits Plant Dictionary

**DOI:** 10.1101/2023.06.16.545047

**Authors:** Elizabeth H. Wenk, Hervé Sauquet, Rachael V. Gallagher, Rowan Brownlee, Carl Boettiger, David Coleman, Sophie Yang, Tony Auld, Russell Barrett, Timothy Brodribb, Brendan Choat, Lily Dun, David Ellsworth, Carl Gosper, Lydia Guja, Gregory J. Jordan, Tom Le Breton, Andrea Leigh, Patricia Lu-Irving, Belinda Medlyn, Rachael Nolan, Mark Ooi, Karen D. Sommerville, Peter Vesk, Mathew White, Ian J. Wright, Daniel S. Falster

## Abstract

Traits with intuitive names, a clear scope and explicit description are essential for all trait databases. Reanalysis of data from a single database, or analyses that integrate data across multiple databases, can only occur if researchers are confident the trait concepts are consistent within and across sources. The lack of a unified, comprehensive resource for plant trait definitions has previously limited the utility of trait databases. Here we describe the AusTraits Plant Dictionary (APD), which extends the trait definitions included in the new trait database AusTraits. The development process of the APD included three steps: review and formalisation of the scope of each trait and the accompanying trait description; addition of trait meta-data; and publication in both human and machine-readable forms. Trait definitions include keywords, references and links to related trait concepts in other databases, and the traits are grouped into a hierarchy for easy searching. As well as improving the usability of AusTraits, the Dictionary will foster the integration of trait data across global and regional plant trait databases.

## Background & Summary

Large-scale analyses of trait data are now commonplace across many scientific disciplines, from vegetation modelling, to evolutionary dynamics and conservation planning ^1, 2^. At the broadest level, a trait is any morphological, physiological, chemical, or life history feature of an organism that can be documented ^3^. Traits capture the enormous heterogeneity in form and function across individuals, populations, or taxonomic units. Variation in trait values reflects the ecological and evolutionary processes that give rise to functional diversity and, in turn, is thus used to define and describe units of biodiversity (e.g., species) ^4^. For vascular plants, the increasing integration of big trait datasets into studies of plant ecology and evolution can be attributed to the rapid growth in databases that collate and/or harmonise collections of field-based observations for re-use ^5^. Some plant trait databases are global ^6–8^, while others have regional ^9–12^ or taxonomic ^13^ scopes. Some target specific organs or functions ^14^, and others are more general, such as floras aimed at plant identification ^15^. Combined with the growing interest in plant traits, the surge in available data is expanding our ability to answer a wide range of questions about the global flora ^6, 16^.

Usefully and accurately capturing the wonderful diversity of plant form and function to address ecological, biogeographic and evolutionary questions involves the non-trivial challenge of reconciling many and often conflicting definitions of plant traits. Garnier^17^ wrote of the “semantic bazaar” in trait ecology, referring to the diversity of possible meanings for a single trait name. For instance, does plant height refer to vegetative height or the height of the highest inflorescence, the height of a typical adult or that of the tallest individual? Is leaf length the length of the leaf blade or does it include petiole length? Without definitions, data cannot be easily reused or merged across trait databases, as the trait names by themselves might not clearly indicate the “trait concept” ^5^. Moreover, as each researcher sees the diversity of form and function in the natural world through a unique lens, the same physical feature on the same plant may be scored as being part of different trait concepts or given a different value of the same trait (Figure 1).

**Figure 1.**
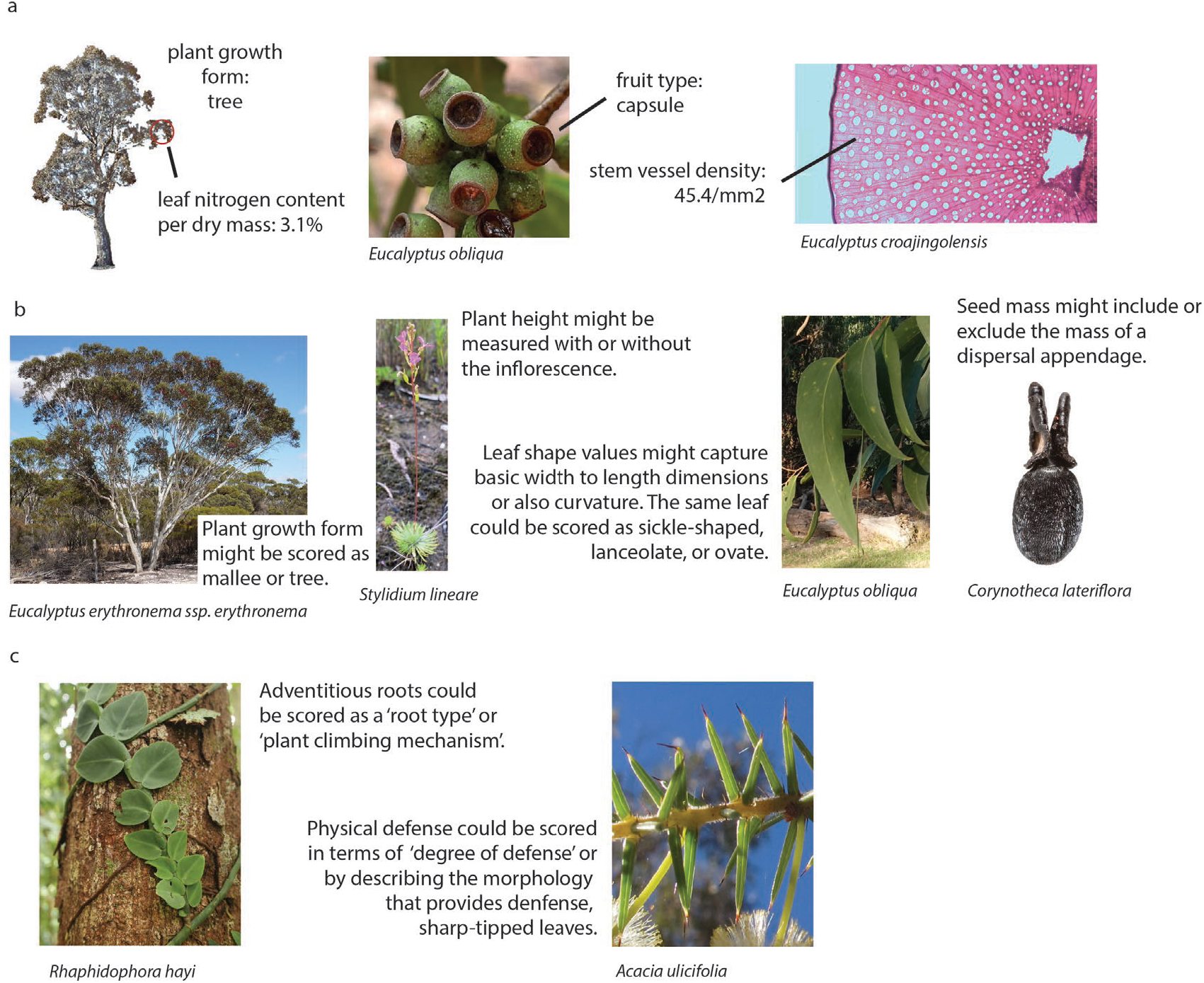
Explicit definitions and value descriptions are needed to reconcile inconsistencies in how researchers align plant phenotypic diversity with particular traits and trait values. a) For some taxa, for some phenotype observations, all researchers are likely to assign the same observation to the same trait and trait value; b) For other taxa, the same trait might not be consistently scored, especially without explicit definitions; c) Some phenotypes will be aligned to different traits or different trait values by different researchers, especially if clear trait and trait value descriptions are not available. Photo credits: Russell Barrett (*Corynotheca lateriflora* seed); John Cull, iNaturalist (*Eucalyptus obliqua* leaves); Gillian Kowalick (*Eucalyptus croajingolensis* cross-section); Dean Nicolle, iNaturalist (*Eucalyptus erythronema* subsp*. erythronema*); Elizabeth Wenk (*Acacia ulicifolia*; *Rhaphidophora hayi*); Dylan Wishart, iNaturalist (*Eucalyptus obliqua* fruits); hughberry, iNaturalist (*Stylidium lineare*).

Ideally, the relationships between different phenotypes and terms would be standardised, allowing researchers to easily reuse data in new contexts. Just as a taxon concept can be described as “a circumscribed set of organisms” (https://github.com/tdwg/tnc/issues/1), a trait concept delimits a collection of trait values pertaining to a distinct characteristic of a specific part of an organism (cell, tissue, organ, or whole organism). Trait names, like taxon names, are associated with each concept, attaching a reusable, interpretable label to each concept, but like taxon names require common terminology across research groups. Currently, research is hindered by the lack of explicit definitions outlining what trait concept a particular trait name refers to, what measurements a specific trait concept encompasses, and the difficulty of reconciling many plausible terms for a single phenotype.

As efforts towards data compilation and database integration have progressed, the need for explicit definitions is increasingly being recognised. Explicit, widely-adopted schemes have long existed for just a few traits (e.g. Raunkiaer’s life forms)^17^ and plant morphology books have long offered a rich vocabulary to describe plant parts ^18, 19^. Meanwhile, trait handbooks have emerged in the ecology and evolutionary biology literature as tools for standardising measurements and terminology ^21–26^. Individual trait databases are also increasingly incorporating explicit trait definitions, enumerating allowable categorical trait values and linking their trait definitions to trait handbooks or published trait ontologies (Table 1)^7, 10, 17, 27–29^. The Thesaurus Of Plant Characteristics (TOP; https://top-thesaurus.org/^17^ was an initiative to define trait concepts for traits in the TRY plant trait database (Table 1) ^6^. Still a work in progress, it was the first extensive effort to publish ecological plant trait definitions that included expected units, allowable categorical trait values and references. There also exist more formal vocabularies put forward by the Open Biological and Biomedical Ontology Foundry (OBO; https://obofoundry.org). One of the OBO Foundry ontologies, the Plant Trait Ontology (PTO; https://bioportal.bioontology.org/ontologies/PTO; Table 1) ^30^, was the first extensive formal ontology of plant traits to be published, including definitions for hundreds of traits relevant to agricultural research organised into an intuitive hierarchy. EnvThes likewise offers a formally published ontology to support Long Term Ecological Research (LTER) data (https://vocabs.lter-europe.net/envthes/; Table 1) ^32^ and is focused on ecological traits. All these pioneering assemblies of trait definitions have advanced global integration of plant trait definitions, but these works remain incomplete relative to the breadth of trait concepts captured by large trait databases.

**Table 1.** Information specified about trait concepts in a selection of trait thesauri, dictionaries, ontologies and databases. (citations: Plant Trait Ontology ^30^; TOP ^17^; EnvThes ^31^; TRY ^6^; GIFT ^7^; BIEN ^8^; LEDA ^27^; BROT 2.0 ^10^

|  | definitions | links to identical trait concepts | specifies units | specifies allowable ranges | specifies and defines allowable trait values | includes references | Machine readable definitions |
| --- | --- | --- | --- | --- | --- | --- | --- |
| Thesaurus, Dictionary or Ontology |  |  |  |  |  |  |  |
| AusTraits Plant Dictionary (APD) | YES | YES | YES | YES | YES | partially | YES |
| Plant Trait Ontology (PTO) | YES | rarely | NO | NO | partially | NO | YES |
| Thesaurus Of Plant Characteristics (TOP) | YES | NO | YES | NO | YES | partially | partially |
| Thesaurus for long term ecological research, monitoring and experiments (EnvThes) | partially | partially | NO | NO | NO | NO | YES |
| Database |  |  |  |  |  |  |  |
| TRY | partially | NO | YES | NO | NO | NO | NO |
| GIFT | NO | NO | YES | NO | YES | NO | NO |
| BIEN | NO | NO | YES | NO | YES | NO | NO |
| LEDA | YES | NO | YES | YES | YES | YES | NO |
| BROT | YES | YES | YES | partially | YES | YES | NO |

Meanwhile, the core components that must be encapsulated to fully define a trait concept have been established by emerging standards in bioinformatic platforms. They include the trait’s name (label), a concise but explicit description, standard units (for numeric traits) and allowable range (for numeric traits) or literal values (for categorical traits) ^29, 32–34^. Additional fields to enhance trait findability include keywords and a trait hierarchy. Interoperability and reusability are increased by including references and links to identical or similar traits in other trait databases. A further step toward making trait definitions FAIR (Findable, Accessible, Interoperable, Reproducible)^35^ is to explicitly link each trait name to a published trait definition ^29^ and to publish a machine-readable trait ontology that accompanies a database or research project.

Despite these multiple efforts from multiple groups, the research community currently lacks comprehensive compilations of definitions that can be readily applied to new data. The existing trait dictionaries, thesauri, and ontologies (Table 1) document an insufficient breadth of traits or offer only partial trait definitions, omitting information such as defined allowable categorical trait values and preferred units, limiting their reuse. For example, we could not find an existing dictionary that contained a definition for a sufficient diversity of traits, with enough detail, or in machine readable format, to support usage of the new AusTraits database. AusTraits is a large continental plant trait database, that currently includes more than 1.25 million data records (v4.1.0) spanning more than 500 traits and nearly all of Australia’s 26,000 plant species ^11^. It includes data for a broad selection of traits including those related to plant morphology, fire ecology, life history, plant physiology, and nutrient contents. The AusTraits workflow requires each trait concept to be linked to allowable trait values, allowable ranges, and accepted units. As the project could not reuse an existing resource, AusTraits developed its own trait dictionary, which is available on the project’s GitHub repository (https://github.com/traitecoevo/austraits.build) and as a data object embedded within the trait database (Figure 2). The traits initially had informal definitions, developed by the AusTraits team, which referenced published trait handbooks, other reference books or an existing thesaurus or glossary, if available, or else were developed in conversation with researchers who contributed data for a unique trait to the database. This process allowed the database to expand rapidly and efficiently, without being limited by availability of dictionaries, whilst still documenting trait definitions, preferred units, numeric ranges and allowable categorical trait values for all traits in the database. While these definitions have allowed AusTraits users to accurately interpret all data within the database and to manually link AusTraits data to those in other trait databases, it was apparent that the utility of AusTraits would be further enhanced by harmonising trait definitions through a formal vocabulary.

**Figure 2.**
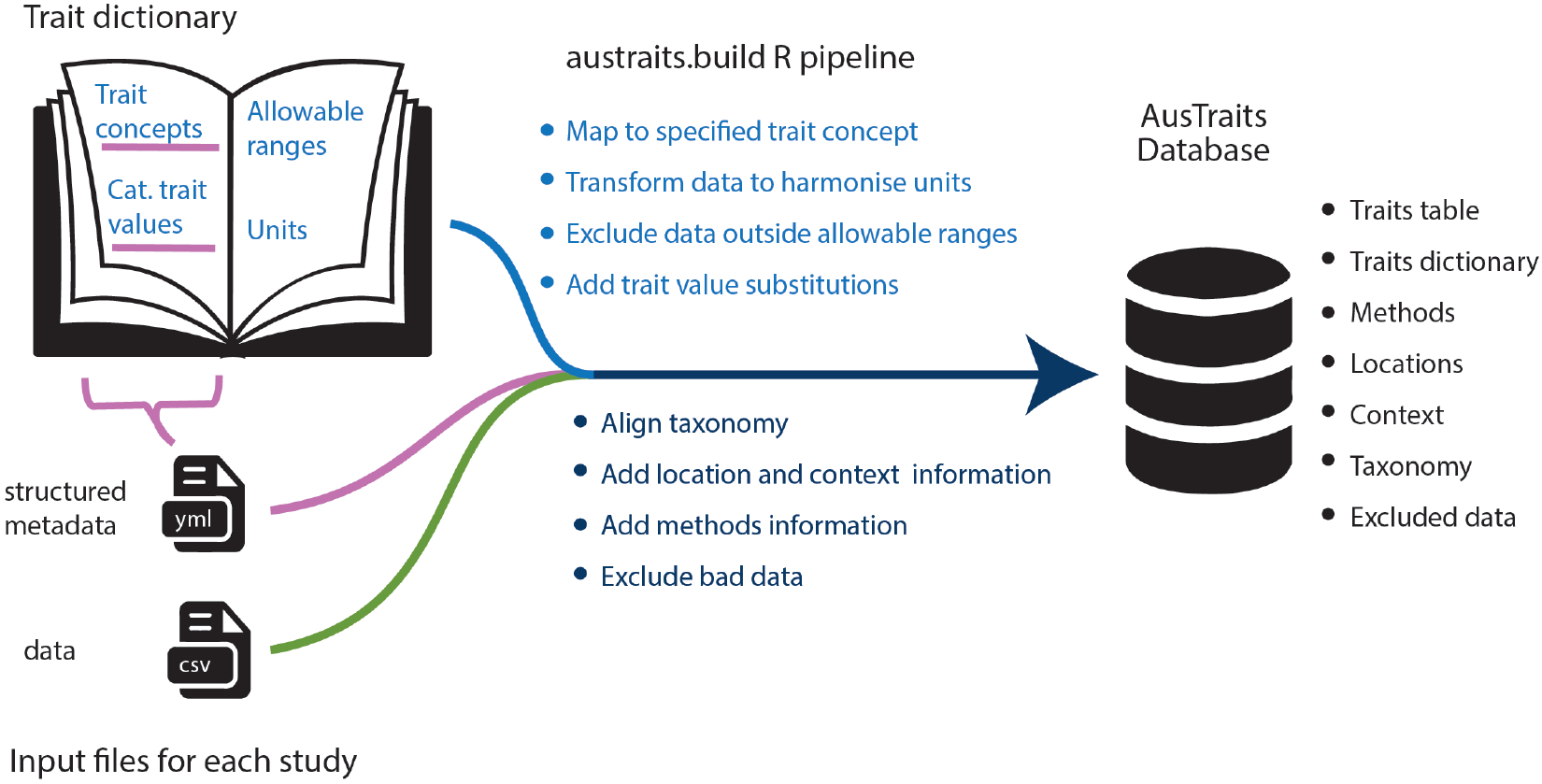
A trait dictionary is an essential component of the AusTraits workflow, specifying i) trait concepts, ii) standard units, iii) allowable categorical trait values and iv) allowable ranges for numeric traits. The structured metadata file that accompanies each dataset explicitly maps data columns to specific trait concepts from the dictionary and includes substitutions to align categorical trait values with those in the dictionary. All four elements of each trait definition are then used by the traits.build R pipeline to integrate the data source into the AusTraits database.

In this paper, we present the AusTraits Plant Dictionary (APD). This comprehensive reference for trait data in AusTraits is also a contribution intended to further integrate global plant trait databases. The APD expands the original AusTraits trait definitions into a formally published dictionary that spans the breadth of trait data included in the database. Most of these definitions will be useful in a global context and expand what is available in existing resources. We used a rigorous review process to refine trait descriptions, added additional metadata to each definition and released the trait dictionary in both a human-friendly and a machine-readable format. Our goal was to progress the global integration of plant trait data in two key ways: first, to create a resource that allowed all data within AusTraits to be effectively mapped onto semantically distinct trait concepts, enhancing the usability of AusTraits; and second, to link this information to other trait databases, thesauri, ontologies and trait handbooks, to both allow the reuse of the APD definitions and facilitate analysis of AusTraits data in combination with data from other databases.

The APD takes the plant trait ecology research community closer to having a global trait dictionary. In addition to supporting AusTraits, we hope that our approach of reviewing and reconciling the often-conflicting trait concepts and descriptions and making them FAIR means we have built a resource that can be reused and built upon by other research initiatives in a global context. Currently, trait concepts and categorical trait values are mostly restricted to traits and terms required to map AusTraits data, but we expect the dictionary to expand over time to support traits and trait values present in other trait databases. For instance, the APD currently has only sparse coverage of root traits, completely lacks traits pertaining to tissue decomposition rates, and is missing some key traits in the hydraulics literature. While a first version of the APD is now available, we expect to continually build upon the dictionary on the project’s GitHub portal, offering successive releases. A customised GitHub issue template allows researchers from across the plant trait research community to suggest additional traits to add to this initiative. A submission would include a proposed trait concept to add, a trait label and description, allowable ranges or values, and references. Once reviewed by the AusTraits team these trait concepts could be included in future releases.

## Methods

There were three components to building a dictionary for these traits. First, we reviewed and revised each trait concept, minimising ambiguity in its scope and writing an explicit, yet concise trait description. Second, we added metadata fields to each trait definition. Third, we compiled all trait concepts into a single resource, output simultaneously in both human and machine-readable formats.

### Reviewing and revising trait concepts, an overview

#### Preliminary review

Through a preliminary review we divided all traits into three groups: 1) trait concepts that were clear and simple and could be reviewed by just the core AusTraits team; 2) trait concepts that required a brief review by experts; and 3) trait concepts where the trait’s scope or allowable values required significant discussion amongst experts; these were reviewed in a series of workshops (Table 2). For the 149 traits that were the content of an element, isotope, metabolite, or other biochemical compound in a specific plant organ, tissue or cell, the meaning and scope of the trait was usually unambiguous and universally agreed upon; few of these traits required a review outside the core AusTraits team. A review by just the core AusTraits team was also sufficient for 134 additional traits with very explicit, simple definitions, or that were trait concepts linked directly to a publication and accompanying dataset.

**Table 2.**
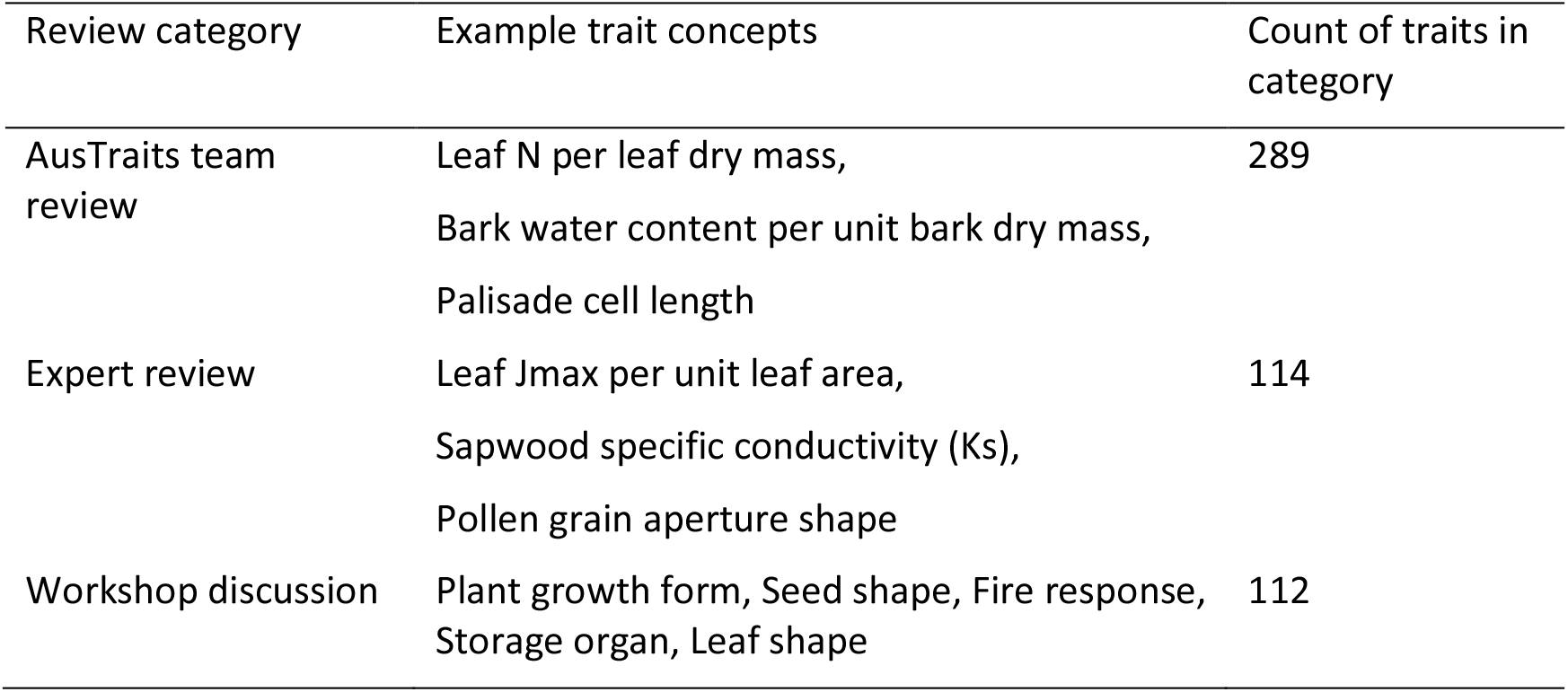
Traits were divided into clusters requiring different styles of review

#### Expert reviews

116 physiological and floral traits were reviewed by experts with extensive experience. These reviewers were able to efficiently identify unrealistic allowable value ranges, nonstandard units, incomplete trait descriptions and call attention to missing or inappropriate key references.

#### Workshop reviews

112 traits were allocated to a specialist workshop, generally because they contained long lists of synonymous or poorly defined categorical trait values or were traits measured differently by different groups of researchers. For traits that required an extensive review, we used a series of small (5–10 person) workshops that brought together researchers who would ideally apply an identical trait concept to diverse research situations. The workshops included researchers at government agencies, universities or herbaria; researchers who were functional ecologists, taxonomists or systematists; and researchers with expertise in diverse plant communities. Three workshops were conducted covering traits within the realms of ‘Seed and dispersal traits’ (October 2021), ‘Leaf and whole plant vegetative traits’ (August 2022) and ‘Fire response and regeneration traits’ (May 2023); each was comprised of 4 or 5 two-hour workshop sessions. Moderated by AusTraits team members, each session was dedicated to clarifying the trait concepts, refining the trait descriptions, identifying key references and carefully compiling a list of allowable trait values that was succinct and distinct. The trait workshops identified trait concepts that were too vague, trait concepts that lacked semantic clarity and curated categorical trait values.

### Completing trait definitions

The core goal of all reviews was to delineate trait concepts and lists of trait values to which all data submitted to AusTraits could be unambiguously mapped. The outcomes of the workshops, expert reviews and internal reviews were used to write a trait description and propagate additional metadata for each trait concept.

#### Trait descriptions and comments

A clear, explicit and comprehensive trait description was drafted for each trait. Whenever possible, trait descriptions were closely aligned to those in trait handbooks, reference books, research papers describing key methods and existing trait ontologies. Following the example set by formal ontologies (e.g. the OBO Foundry ontologies PATO, PO and PTO) and the TOP Thesaurus, a second formal description was drafted for each trait where all technical terms were linked to classes (words; concepts) in published ontologies. This removed ambiguity in what was meant by a term and required that all definitions were written with reference to a narrow list of words. Although this formal method generated a unique definition for each trait, the less formal, non-annotated trait descriptions were considered essential to convey the trait concepts to users, as the encoded descriptions are often awkward to read and interpret. In addition, a comments field provided a location to document notes, including referencing similar traits, best practice measurement methods, important context variables and possible sources of error within the amalgamated data. For instance, the definition for the trait “leaf area” could include a comment indicating that although only leaf area data are meant to be mapped to this trait concept, many authors will merge leaflet and leaf area data under the title “leaf area” and therefore trait databases, such that AusTraits, will contain a mix of leaf and leaflet area data under the “leaf area” trait. Meanwhile, for photosynthetic rate traits the comments field could indicate that it is best practise to document leaf temperature as a context property.

In addition to the trait description and comments, descriptive metadata fields were added to each trait concept (Table 3) ^36^. The metadata fields include those required for data processing (e.g. allowable ranges and allowable trait values), those that increase trait concept findability (e.g. keywords), those that properly document the source and scope of the trait (e.g. references, reviewers) and those that increase trait concept interoperability across datasets (e.g. matches to other databases).

**Table 3.**
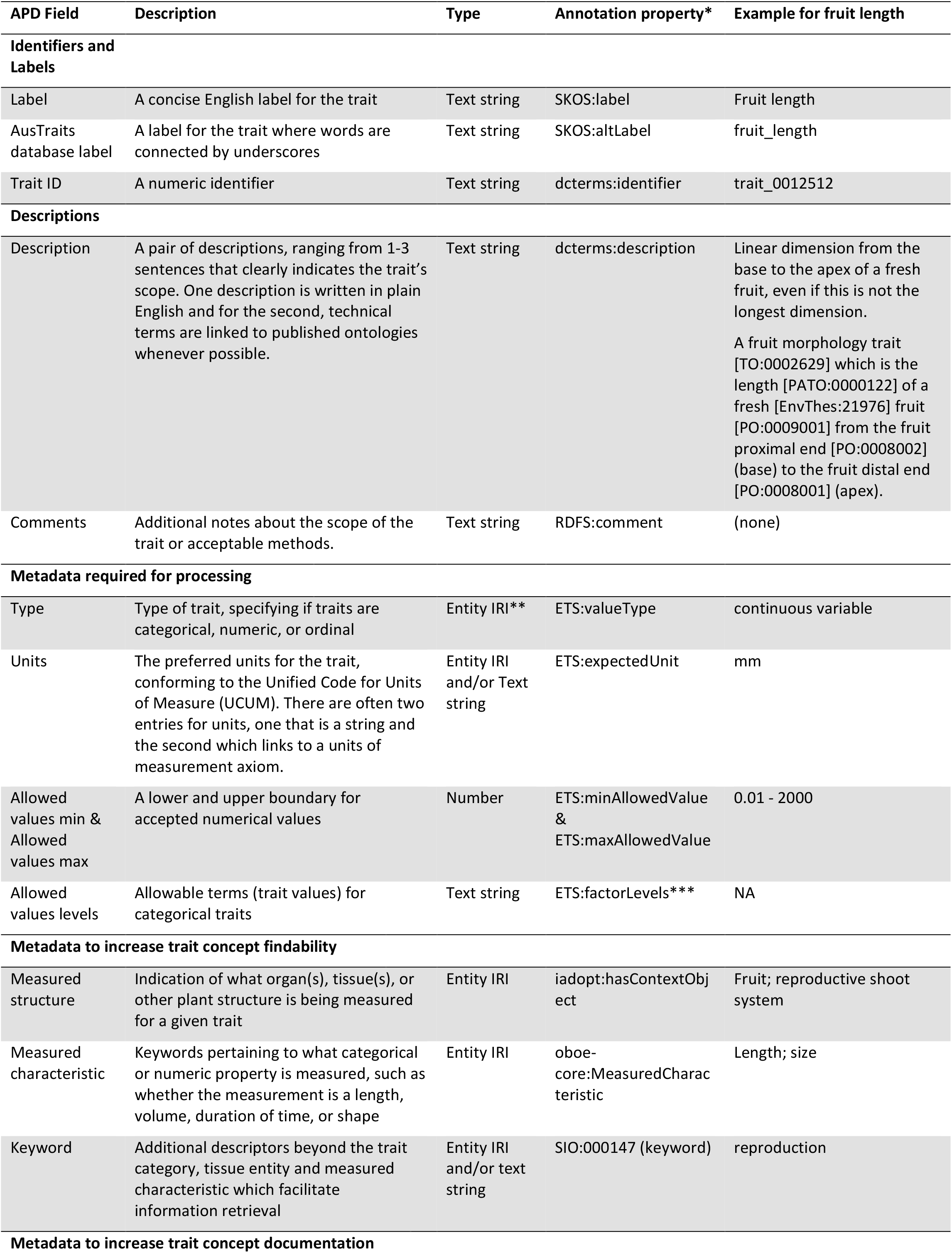
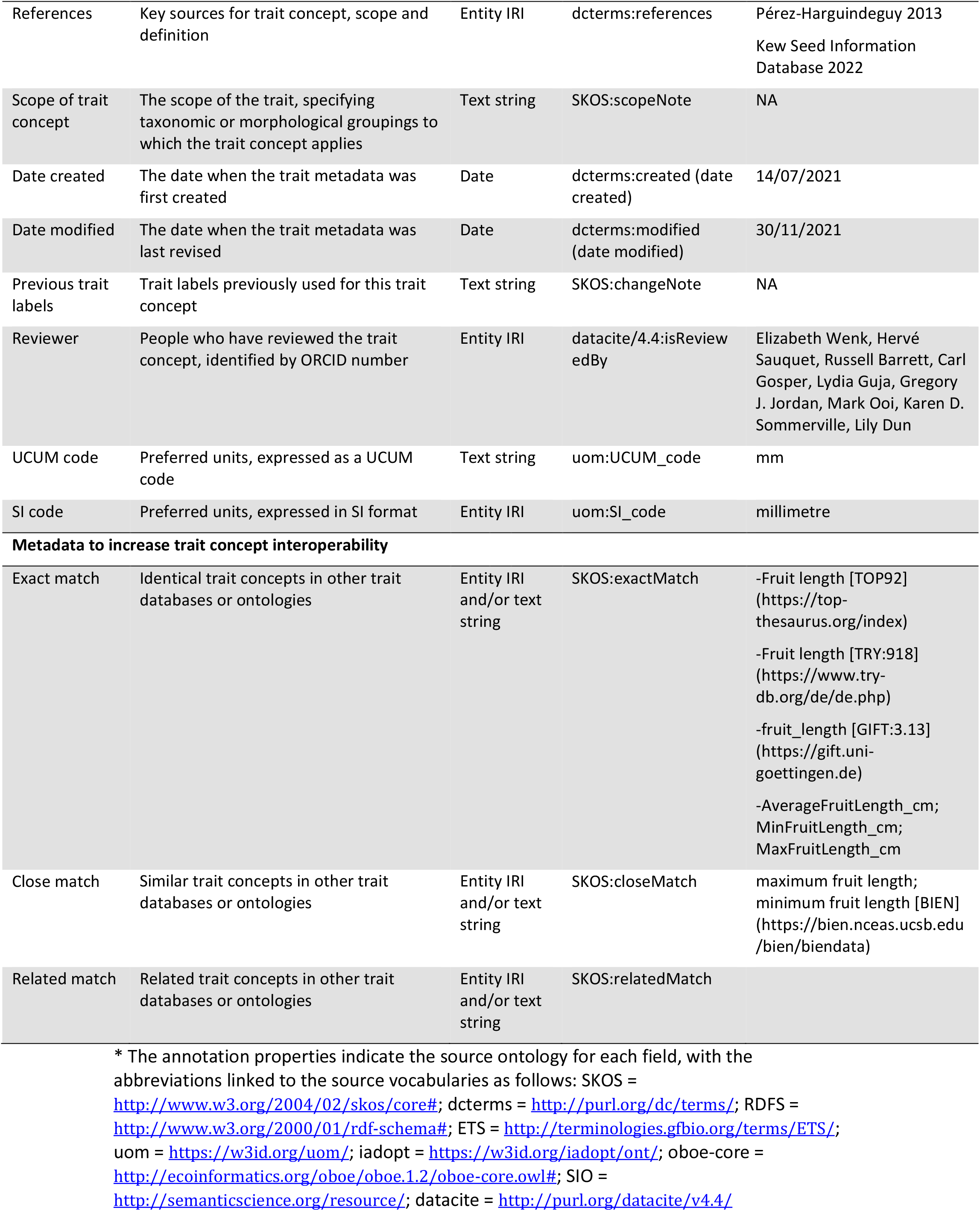
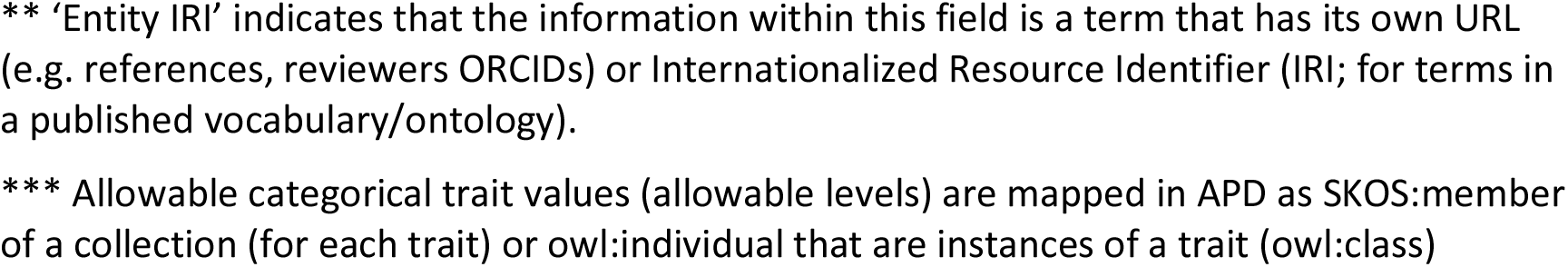
Metadata provided for each trait concept, including a description of each metadata field and the published annotation property onto which this information is mapped.

| APD Field | Description | Type | Annotation property* | Example for fruit length |
| --- | --- | --- | --- | --- |
| **Identifiers and Labels** |  |  |  |  |
| Label | A concise English label for the trait | Text string | SKOS:label | Fruit length |
| AusTraits database label | A label for the trait where words are connected by underscores | Text string | SKOS:altLabel | fruit_length |
| Trait ID | A numeric identifier | Text string | dcterms:identifier | trait_0012512 |
| **Descriptions** |  |  |  |  |
| Description | A pair of descriptions, ranging from 1-3 sentences that clearly indicates the trait's scope. One description is written in plain English and for the second, technical terms are linked to published ontologies whenever possible. | Text string | dcterms:description | Linear dimension from the base to the apex of a fresh fruit, even if this is not the longest dimension.<br>A fruit morphology trait [TO:0002629] which is the length [PATO:0000122] of a fresh [EnvThes:21976] fruit [PO:0009001] from the fruit proximal end [PO:0008002] (base) to the fruit distal end [PO:0008001] (apex). |
| Comments | Additional notes about the scope of the trait or acceptable methods. | Text string | RDFS:comment | (none) |
| **Metadata required for processing** |  |  |  |  |
| Type | Type of trait, specifying if traits are categorical, numeric, or ordinal | Entity IRI** | ETS:valueType | continuous variable |
| Units | The preferred units for the trait, conforming to the Unified Code for Units of Measure (UCUM). There are often two entries for units, one that is a string and the second which links to a units of measurement axiom. | Entity IRI and/or Text string | ETS:expectedUnit | mm |
| Allowed values min & Allowed values max | A lower and upper boundary for accepted numerical values | Number | ETS:minAllowedValue & ETS:maxAllowedValue | 0.01 - 2000 |
| Allowed values levels | Allowable terms (trait values) for categorical traits | Text string | ETS:factorLevels*** | NA |
| **Metadata to increase trait concept findability** |  |  |  |  |
| Measured structure | Indication of what organ(s), tissue(s), or other plant structure is being measured for a given trait | Entity IRI | iadopt:hasContextObject | Fruit; reproductive shoot system |
| Measured characteristic | Keywords pertaining to what categorical or numeric property is measured, such as whether the measurement is a length, volume, duration of time, or shape | Entity IRI | oboe-core:MeasuredCharacteristic | Length; size |
| Keyword | Additional descriptors beyond the trait category, tissue entity and measured characteristic which facilitate information retrieval | Entity IRI and/or text string | SIO:000147 (keyword) | reproduction |
| **Metadata to increase trait concept documentation** |  |  |  |  |
| References | Key sources for trait concept, scope and definition | Entity IRI | dcterms:references | Pérez-Harguindeguy 2013<br>Kew Seed Information Database 2022 |
| Scope of trait concept | The scope of the trait, specifying taxonomic or morphological groupings to which the trait concept applies | Text string | SKOS:scopeNote | NA |
| Date created | The date when the trait metadata was first created | Date | dcterms:created (date created) | 14/07/2021 |
| Date modified | The date when the trait metadata was last revised | Date | dcterms:modified (date modified) | 30/11/2021 |
| Previous trait labels | Trait labels previously used for this trait concept | Text string | SKOS:changeNote | NA |
| Reviewer | People who have reviewed the trait concept, identified by ORCID number | Entity IRI | datacite/4.4:isReviewedBy | Elizabeth Wenk, Hervé Sauquet, Russell Barrett, Carl Gosper, Lydia Guja, Gregory J. Jordan, Mark Ooi, Karen D. Sommerville, Lily Dun |
| UCUM code | Preferred units, expressed as a UCUM code | Text string | uom:UCUM_code | mm |
| SI code | Preferred units, expressed in SI format | Entity IRI | uom:SI_code | millimetre |
| Metadata to increase trait concept interoperability |  |  |  |  |
| Exact match | Identical trait concepts in other trait databases or ontologies | Entity IRI and/or text string | SKOS:exactMatch | -Fruit length [TOP92] (https://top-thesaurus.org/index)<br>-Fruit length [TRY:918] (https://www.try-db.org/de/de.php)<br>-fruit_length [GIFT:3.13] (https://gift.uni-goettingen.de)<br>-AverageFruitLength_cm;<br>MinFruitLength_cm;<br>MaxFruitLength_cm |
| Close match | Similar trait concepts in other trait databases or ontologies | Entity IRI and/or text string | SKOS:closeMatch | maximum fruit length;<br>minimum fruit length [BIEN] (https://bien.nceas.ucsb.edu/bien/biendata) |
| Related match | Related trait concepts in other trait databases or ontologies | Entity IRI and/or text string | SKOS:relatedMatch |  |

\* The annotation properties indicate the source ontology for each field, with the abbreviations linked to the source vocabularies as follows: SKOS = http://www.w3.org/2004/02/skos/core#; dcterms = http://purl.org/dc/terms/; RDFS = http://www.w3.org/2000/01/rdf-schema#; ETS = http://terminologies.gfbio.org/terms/ETS/; uom = https://w3id.org/uom/; iadopt = https://w3id.org/iadopt/ont/; oboe-core = http://ecoinformatics.org/oboe/oboe.1.2/oboe-core.owl#; SIO = http://semanticscience.org/resource/; datacite = http://purl.org/datacite/v4.4/
\*\* 'Entity IRI' indicates that the information within this field is a term that has its own URL (e.g. references, reviewers ORCIDs) or Internationalized Resource Identifier (IRI; for terms in a published vocabulary/ontology). \*\*\* Allowable categorical trait values (allowable levels) are mapped in APD as SKOS:member of a collection (for each trait) or owl:individual that are instances of a trait (owl:class)

#### Metadata required for processing

The R pipeline that compiles AusTraits requires five pieces of information for each trait concept: the label (i.e., a trait name), the trait type (numeric versus categorical), the allowable range of values for numeric traits, the standardised units for numeric traits, and the allowable trait values for categorical traits (Figure 2; Table 3).

#### Metadata to increase trait concept findability

Metadata should also include descriptors that aid in the discovery of the resource, here the individual trait concepts. In addition to offering a trait hierarchy, APD includes three fields to increase trait findability: the plant structure being measured (e.g., stem, leaf, root, whole plant, flower), the characteristic being measured (e.g., mass, shape, force) and additional keywords (Table 3).

#### Metadata to increase trait concept documentation

Each trait in the APD includes metadata to record the date trait concepts were described and revised; the people involved in reviewing each trait concept and trait definition; its applicability (i.e., scope; the trait might only be scorable for specific taxonomic groups or in plants that have leaves); and past labels (names) used for the identical trait concept. In addition, references were linked to traits whenever possible; these included trait handbooks describing the trait, manuscripts introducing or championing the trait and review papers noting the best traits to measure to document a particular plant function. Two fields link the standardised units to published vocabularies, described below.

#### Metadata to increase trait concept interoperability

A cluster of metadata elements promote the interoperability of this resource with other databases by documenting trait concepts in other trait databases or ontologies that are identical, similar, or related to a specific trait concept in the APD (Table 3). Trait concepts from TRY ^6^, TOP ^17^, GIFT ^7^, LEDA ^27^, BIEN ^8^, BROT 2.0 ^10^, the Palm Traits Database ^13^, the Plant Trait Ontology ^30^ and EnvThes ^31^ were cross-mapped to trait concepts in the APD.

#### Mapping metadata fields to concepts in published vocabularies

In a standard tabular format, the metadata fields would be the column headers, each specifying a different piece of information documented about the trait. In a formal ontology, each metadata field included in the APD must be matched to an appropriate annotation property. These are published, formally defined terms for ‘label’, ‘description’, etc. (Table 3). By preference, metadata fields are linked to concepts defined by the often-used Simple Knowledge Organization System (SKOS; https://www.w3.org/TR/skos-primer/^37^, Resource Description Framework (RDF; https://www.w3.org/TR//rdf11-primer/^38^, or dcterms (Dublin Core Metadata Initiative) ^39^ vocabularies. Properties defined by the Ecological Trait-data Standard (ETS) ^32^ were also reused; this schema is establishing itself as a well-designed ecological trait database structure. Units were aligned to the Unified Code for Units of Measure (UCUM; https://ucum.org/ucum) standard with specific machine-readable representations of each unit downloaded from the Units of Measurement (UOM) portal (https://units-of-measurement.org/). The UCUM standard follows clear, simple rules, but also has a flexible syntax for documenting notes that are recorded as part of the ‘unit’ for specific traits, yet are not formally units, in curly brackets ^40^. For instance, {count}/mm2 or umol{CO2}/m2/s, where the actual units are 1/mm2 and umol/m2/s. An added advantage is that the UOM representations include links to identical units in a collection of other published ontologies. Properties not present in any of these ontologies were mapped to ones in the Semantic Science Information Ontology (SIO) ^41^, the Extensible Observation Ontology (OBOE) ^42^, Datacite ^43^ and the iAdopt Ontology ^44^ (Table 3).

Within each APD trait concept, some trait metadata fields were simply text strings (i.e., trait description), while other metadata values were themselves published concepts with a Uniform Resource Identifier (URI) (an inclusive term, that encompasses both URLs and Internationalized Resource Identifiers [IRIs]). For instance, references mostly have a DOI (digital object identifier), reviewers were identified by their ORCID numbers (Open Researcher and Contributor Identifier) and keywords were all identified by their URI’s from various published ontologies.

### Trait hierarchy

For ease of grouping trait concepts, we established a trait hierarchy into which the traits could be slotted. At the highest level, all traits within the APD could be divided into four categories: biochemical traits, morphological traits, physiological traits and life history traits (Table S1). Three of these were exact matches to classes defined by the Plant Trait Ontology (plant biochemical trait (http://purl.obolibrary.org/obo/PTO_0000277); plant morphology trait (http://purl.obolibrary.org/obo/PTO_0000017); and biological process trait, physiological process trait (http://purl.obolibrary.org/obo/PTO_0000283)), while life history trait was defined within APD. Additional hierarchical levels were established, again using a combination of terms from the Plant Trait Ontology and ones defined within APD (Table S1). The trait hierarchy was mapped into the formal ontology as nested SubClasses, cascading down from the top concept, a Plant Trait (http://purl.obolibrary.org/obo/PTO_0000000).

### Building APD into a machine-readable resource

The primary output for the trait concepts and their associated metadata needed to be in a machine-readable format that could both be stored on the project’s GitHub repository and published online through the Australian Research Data Common’s (ARDC) Research Vocabularies Australia (https://ardc.edu.au/services/research-vocabularies-australia/). Each term defined within the APD requires a unique and stable URI. This includes not just the trait concepts, but also the allowable categorical trait values, the trait groupings within the trait hierarchy, and the selection of terms within a glossary. Although the APD outputs are archived on the project’s GitHub repository, we chose to register the APD namespace with the URI redirection service w3id.org to ensure the permanency of the URI’s even if our project repository were to be moved. The trait concepts, trait groupings for the hierarchy, and allowable categorical trait values are within one schema, w3id.org/APD/traits/ while the glossary terms are in a second schema, w3id.org/APD/glossary.

A machine-readable representation was built using an R script that first merged seven separate data tables into a single table formatted as RDF Triples, the core unit of the Resource Description Framework (RDF) data model. With the triples format, all information content is collapsed into a single long-format document with three columns, the subject, the predicate, and the object. The subject is always the URI for a concept or term, and, for the APD, included both the URI’s within the w3id.org/APD namespace as well as concepts within the ancillary tables, such as ORCIDs for reviewers, DOI’s for references, or URI’s for concepts reused from published vocabularies. The predicate indicates a property of the object that can be described. The predicates in AusTraits are the annotation properties in Table 3 and additional terms specified under ‘Column’ in Tables S2-S10. Each predicate is also a URI. The object is the value for the specific predicate for the specific object.

Spreadsheets with data that were converted to triples include: 1) the core trait concepts and their metadata; 2) trait references; 3) trait reviewers (by ORCID); 4) classes from published ontologies; 5) terms from the Units of Measurement Ontology; 6) allowable categorical trait values; 7) the trait hierarchy under which the traits in the APD could be grouped. Terms sourced from published ontologies or other sources were mapped into APD as their own entities to ensure their labels and descriptions were included within APD, rather than simply being identified by an URI. As each value of a property of an object is in a new row in triples format, there may be more than 30 rows of data for a single trait concept (Table S2), and, in total, there are more than 33,000 rows of unique object-predicate-value combinations within the APD.

The R package rdflib ^45^ was used to serialise the table of triples into RDF objects, output in Turtle (APD.ttl), N-Triple (APD.nt), N-Quad (APD.nq) and JSON Linked Data (APD.json) formats.

The RDF serialisations were complemented by two derivatives, created from the N-Triple output using a combination of R and Quarto scripts (Figure 4). The first is a HTML landing page for human interaction with the machine-readable formats (https://traitecoevo.github.io/APD/index.html) to which all searches for individual concept URIs are automatically redirected. And second is the YAML (.yml) file required by the AusTraits workflow to compile the database. It includes only the trait labels, trait description, type, allowable range, allowable trait values and required units and is located within the austraits.build GitHub repository (https://github.com/traitecoevo/austraits.build). The YAML format offers a flexible data serialisation format to capture diverse metadata in a single file, as it has a nested format which allows different numbers of levels beneath each header. This permits both easy data input and human interpretation.

**Figure 3.**
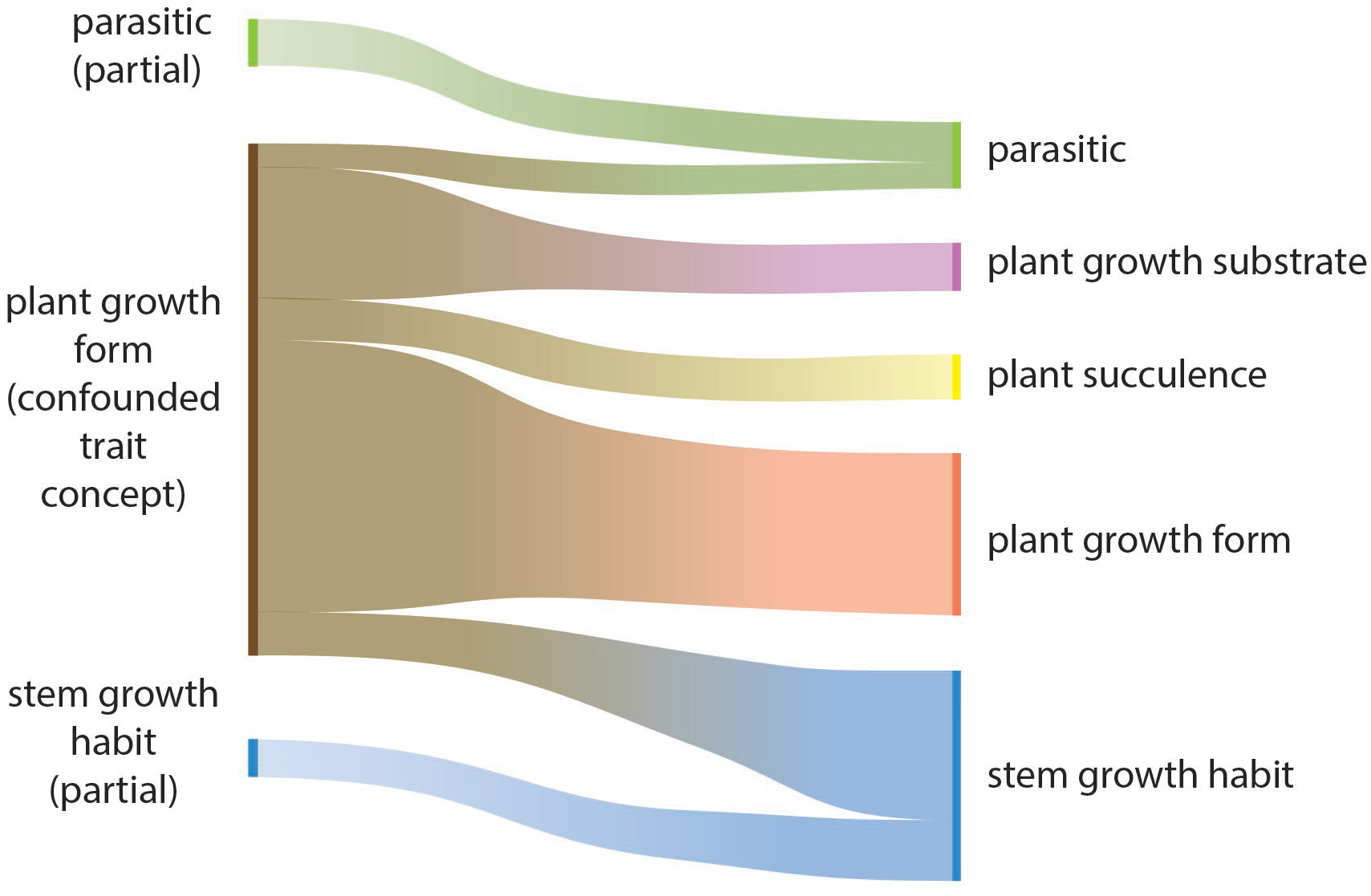
The initial list of trait values mapped to the semantically messy trait concept ‘plant growth form’ and the trait concepts ‘parasitic’ and ‘stem growth habit’ were able to be condensed from 68 to 53, despite adding more detailed trait values to parasitic, plant succulence and stem growth habit traits. For the APD, the retained trait values were mapped across 5 traits: plant growth form, plant succulence, plant growth substrate, parasitic and stem growth habit. The mixing of semantic concepts within ‘plant growth form’ had previously resulted in hybrid terms which could now be eliminated, such as ,shrub_aquatic-.

**Figure 4.**
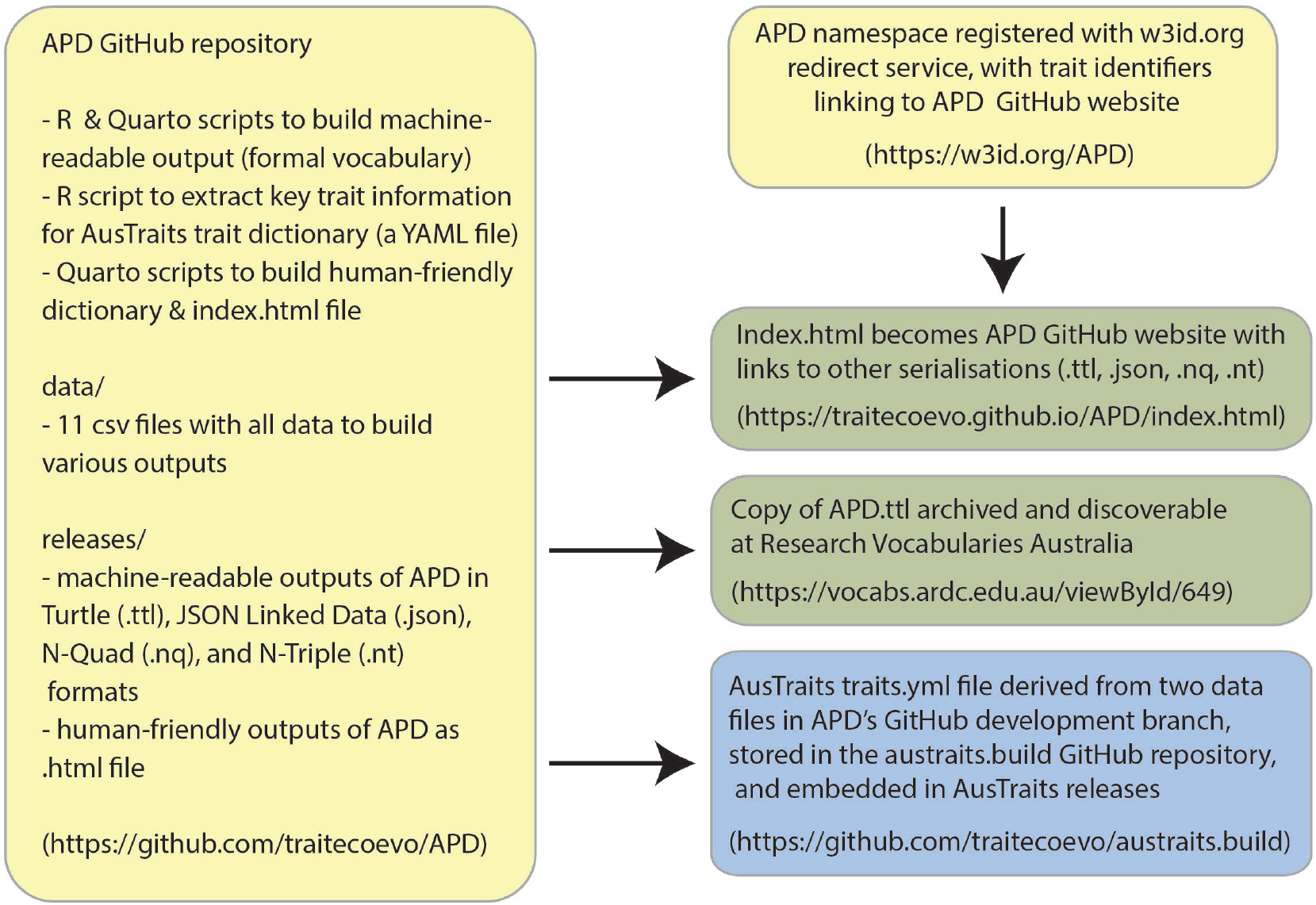
The APD inputs and stored on the project GitHub repository, the versioned outputs archived on the GitHub repository, Zenodo, and at the Australian Data Research Common’s Research Vocabulary Australia (RVA) portal. APD has been registered as a namespace within w3id.org, with term URI’s redirecting back to an HTML landing page within the GitHub repository. The APD inputs are also used to generate the traits.yml file required to build the AusTraits trait database.

## Data Records

### Trait concepts and allowable trait values

In total, APD includes 515 traits, including 112 categorical traits and 403 numeric traits (Table S3). These vary from well-known traits like leaf area to bespoke ones like leaf pendulousness that are measured only for specific research questions. The internal reviews, expert reviews and reviews through the trait workshops all worked toward clarifying trait concepts and developing clear trait descriptions and appropriate lists of allowable categorical trait values.

#### Trait concept, label and description too vague

The vocabulary workshops uncovered several instances where trait names were ambiguous and may have led to the misinterpretation of data. For instance, the trait ‘leaf angle’ was defined as the angle between the stem and the leaf blade, but it was identified that the data in AusTraits referred to the leaf blade’s angle relative to the solar zenith. There are now two traits in the APD with more explicit labels and definitions, leaf axil angle and leaf inclination angle. Another example of a semantically unclear trait label was the trait capturing the hairiness of juvenile leaves. It was unclear if these were the leaves on a juvenile plant or the juvenile (regrowth) leaves on an adult plant following disturbance. Again, it was necessary to adopt two separate traits whose scopes were more explicit. In addition, by linking the terms in the trait description to ontologies, it was possible to clearly distinguish between a leaf on a ‘juvenile plant’ (https://purl.obolibrary.org/obo/PATO_0001190) versus a ‘juvenile leaf’ (https://purl.obolibrary.org/obo/PO_0006339) on an adult plant.

#### Trait concept too broad

There were several traits that were identified as being too broad and including two (or more) semantically distinct concepts; these traits were split into multiple traits with a narrower, explicitly defined scope. For instance, fruit type included both true, botanical fruit types and terms that simply indicated whether a dispersal unit was dry or fleshy. The data initially merged together under fruit type were split into a trait that captured true botanical fruit types, such as achenes and drupes ^46^ and then two traits that indicated specific functions of the fruit, independent of its formal classification, i.e., fruit fleshiness and fruit dehiscence. Plant growth form included terms that pertained not only to the actual entire plant form, but also values indicating whether it was terrestrial, aquatic, or epiphytic and whether it was a parasite. The initial scope of data mapped to plant growth form was divided into a simpler plant growth form which was focused on the plant’s perennating 3-dimensional shape, with ancillary information mapped to plant growth substrate, plant succulence and, in part, a revised stem growth habit and parasitic traits (Figure 3). Trait concepts that are too broad are a global problem and other trait databases have also recently taken the approach of splitting plant growth form into more tractable traits with a clearly defined ‘entity’ and scope ^7, 47^. For the APD, this allowed a considerable reduction in repetitive trait values, such as remapping ‘aquatic_herbs’; ‘aquatic_shrubs’ and ‘aquatic_trees’ to ‘herbs’, ‘shrubs’ and ‘trees’ under plant growth form and as ‘aquatic’ under plant growth substrate.

#### Curating categorical trait values

Certain categorical traits were identified as those most requiring standardisation of trait values and were selected to review during the workshops. These included seed shape, fruit type, dispersal syndrome, leaf shape, leaf type and plant growth form. These were traits for which there were data in many datasets, but which lacked universally agreed upon allowable trait values. Despite attempts to condense terms and align meanings, AusTraits had 50–80 trait values for leaf shape; many were clearly synonymous terms or terms not actually related to the shape of the leaf blade. There were two core reasons for these long lists of terms: 1) traits that integrated data from both the ecology and the systematics communities, with different researchers favouring different sets of terms; and 2) the lack of available vocabulary to describe particular trait phenotypes.

Plant morphologists and taxonomists are equipped with botanical glossaries^19, 20, 48, 49^, offering a detailed vocabulary to describe all nuances of a plant’s morphology. In contrast, while ecologists use these morphological terms when appropriate, ecology datasets also include terms that capture specific functional roles, often using a merging of formal and informal terms. By curating categorical trait values, two core revisions were made. The first was to condense the extensive list of terms in botanical glossaries. Although many researchers in these fields take advantage of this rich descriptive vocabulary, they were amenable to reducing the list of terms allowed as values for a given trait, realising that the fine-grained distinctions were unlikely to have functional significance, but also that many terms were so similar they were unlikely to be used consistently, even by the experts. This concurs with recent research that suggests that all people, even expert botanists, were more likely to correctly identify a plant’s character when there were fewer options to choose from ^50^. Synonymous terms were listed within the description of each trait value (Table 3), clarifying the scope of each trait value retained and facilitating searches for terms that were omitted. For example, for seed surface texture, the final list included 11 trait values, but an additional 28 terms were mapped as synonyms (Table 4). For some traits, appropriate lists of terms were discovered through literature searches or emerged through workshop discussions. For instance, the many leaf shape values could easily be mapped to the terms in a resource established by the Systematics Association Committee 60 years ago^51^, which was not known to most but familiar to one workshop participant.

**Table 4.** Explicitly listing synonyms as part of trait value definitions ensured alternative terminology can be consistently mapped to the term used in the APD, as illustrated here for seed surface texture.

| Trait Value | Synonyms |
| --- | --- |
| bumpy | colliculate, verrucate, papillate, tuberculate, undulate |
| grooved |  |
| netted | reticulate, honey-combed |
| papery | chartaceous |
| pitted | foveolate, foveate, dimpled, lacunose, punctate |
| ribbed | carinate, costate, fluted, lineate, lineolate, ridged, scalariform, striate, strigose |
| rough | scabrous |
| scaly | scurfy, squarrose |
| smooth | glabrous |
| spiny | echinulate |
| wrinkled | rugose, rugulose, bullate |

Some challenges emerged when selecting a list of allowable words to describe the ecological or functional trait values where no succinct, unified list of terms exists. The difficulty is exemplified by plant growth form, where even successive versions of the same trait handbook presented barely overlapping lists of allowable growth form terms^21, 22^, despite this being one of the most recorded traits worldwide. These resources and many others share the use of ‘tree’, ‘shrub’ and ‘herb’, but beyond these terms resources diverge in their list of allowable plant growth form values. Our list uses terms from both of these references as well as many others ^7, 17, 47^, compiling a list onto which all existing AusTraits data could be mapped. Our goal was to balance having enough terms to capture morphological and functional diversity, while allowing for comparative analyses across groupings. For plant growth form, as for other traits, this list included terms of ecological or descriptive significance that might be used only for specific taxa or ecological situations, yet were required for trait measurements in those circumstances. For the Australian flora, terms like ‘mallee’ and ‘hummock’ were deemed essential to describe distinct plant growth forms, although these terms are absent or rarely used globally. In the final list there was a clear scope and description attached to each trait value.

## Data files

The APD GitHub repository (https://github.com/traitecoevo/APD) includes the eleven spreadsheets required to compile the final resource.

**APD_traits.csv** is the core data table, which includes trait labels, trait descriptions and all associated metadata for each trait concept (Table S4). As indicated in Table 3, some columns are textual strings, others are numeric and some refer to pre-existing entities (concepts, classes). The pre-existing entities are documented in an additional four data tables, APD_references.csv, APD_reviewers.csv, APD_units.csv and published_classes.csv.

**APD_references.csv** links each reference indicated in APD_traits.csv to its DOI (or alternative identifier), also providing a title and complete reference (as a string) (Table S5).

**APD_reviewers.csv** links each reviewer indicated in APD_traits.csv to their ORCID number (Table S6).

**APD_units.csv** links the standardised units indicated in APD_traits.csv to their respective URLs in the Units of Measurement Ontology (Table S7). The data table includes a description of the unit, links to its SI and UCUM representation and indicates other ontologies with definitions for this unit.

**published_classes.csv** documents terms from published ontologies used as keywords, measured characteristics, measured structures, or to describe the trait type. The label, description, IRI, scheme URI and scheme prefix are provided for each term (Table S8).

**APD_categorical_values.csv** contains the allowable trait values for each categorical trait, including descriptions of each term and indicating the trait concept to which the term is linked (Table S9).

A challenge in the compilation of APD was that ontologies allow only a single instance of each word to be used, with a single definition. While each trait name is unique, the same term (word) can be used as a categorical trait value for multiple traits with subtly different meanings and possibly different meanings to a pre-existing ontology. Generalising the definitions to be applicable to all instances of its use would mean that its definition would be far broader than implied as a specific trait value for a single categorical trait. The solution for APD was for official trait values to be the merging of the trait label and the term, while the label for the term could be a simple word that might be reused. For instance, the trait value *hairy* is used for five separate traits and for the trait *Juvenile phase leaf hairiness* the formal trait value becomes *leaf_hairs_juvenile_leaves_hairy*.

**APD_trait_hierarchy.csv** indicates the hierarchical structure into which the trait concepts are mapped (Table S10).

**APD_glossary.csv** includes a collection of terms used repeatedly within APD trait concept descriptions or as keywords, but which lacked an appropriate published definition (Table S11).

**APD_annotation_properties.csv** indicates the source, label and description for each of the annotation properties (Table 3) used to capture metadata for the trait concept s (Table S12).

**APD_namespace_declaration.csv** indicates the URI for each vocabulary prefix referenced in APD_traits.csv and serves as the namespace declaration when compiling the RDF representation (Table S13).

**APD_resource.csv** is already in Triples format and includes annotation properties about the core APD resources, APD/traits and APD/glossary (Table S14).

## Access

The data are available under a CC-BY 4.0 license, allowing reuse with attribution. The versioned releases are archived on Zenodo (https://doi.org/10.5281/zenodo.8040789). The version controlled machine-readable Turtle representation is also published through Research Vocabularies Australia, part of the national research infrastructure operated by the Australian Research Data Commons (ARDC) (https://vocabs.ardc.edu.au/viewById/649). The APD GitHub repository (https://github.com/traitecoevo/APD) has both versioned releases and ongoing development versions. The APD namespace (w3id.org/APD) and trait concept URI’s (e.g. https://w3id.org/APD/traits/trait_0000014) also redirect to the versioned releases on the APD GitHub repository.

## Technical Validation

The APD.ttl file (Turtle serialisation) was run through a skos validator to confirm that all relationships were consistent, all URI’s were unique, and that all concepts has labels. The APD.csv (in Triples format) was used to recompile the HTML landing page. The APD_traits.csv and APD_categorical_values.csv files were used to recompile the YAML file for the AusTraits workflow. Deriving the HTML output from the Turtle serialization further and confirming AusTraits continued to build properly from the automatically regenerated YAML file, confirmed the files were complete and the process was accurate.

## Code Availability

The code to compile the data into the selected output formats is available on the APD GitHub repository (https://github.com/traitecoevo/APD).

## Acknowledgements

Input from and conversations with the following people enhanced the review of trait definitions: Will Cornwell, Félix de Tombeur, Saskia Grootemaat, Gillian Kowalick, Thomas Mesaglio, Ruby Stephens and Isaac Towers. We are grateful to Simon Cox, Jon Smillie, Kerry Levett, Melanie Barlow, and Catherine Brady for useful conversations about how to publish a formal vocabulary. We also thank all AusTraits data contributors for providing the data that allowed AusTraits to grow into a sufficiently large, diverse database that the APD was able to emerge as a standalone resource. The AusTraits project received investment (https://doi.org/10.47486/TD044, https://doi.org/10.47486/DP720) from the Australian Research Data Commons (ARDC). The ARDC is funded by the National Collaborative Research Infrastructure Strategy (NCRIS).

## Author contributions

E.H.W., H.S., R.V.G., and D.S.F. conceived the original idea.

E.H.W. led the writing of the manuscript.

E.H.W, R.B, C.B. and D.S.F. led the coding and the technical development of the RDF serialisations.

E.H.W, S.Y. and D.C let the AusTraits team review of trait concepts. T.B., B.C., D.E., B.M, and H.S. offered expert reviews of traits.

E.H.W., H.S., R.V.G, T.A., R.L.B., D.C., L.D., C.G., L.G., G.J.J., A.L., P.L., T.L., R.N., M.O., K.S., P.V.,

I.J.W., M.W., and S.Y. participated in the workshop reviews.

## Competing interests

The authors declare no competing interests.

## Supplementary Tables

**Supplementary Table 1.** Number of traits in each of the hierarchical trait groupings.

| trait grouping | number of<br>trait concepts<br>within<br>group* |
| --- | --- |
| **biochemical trait** |  |
| **mineral and ion content trait** |  |
| leaf mineral and ion content trait |  |
| live leaf mineral and ion content trait | 33 |
| senesced leaf mineral and ion content trait | 16 |
| stem mineral and ion content trait | 17 |
| wood mineral and ion content trait | 7 |
| senesced wood mineral and ion content trait | 6 |
| bark mineral and ion content trait | 9 |
| root mineral and ion content trait | 3 |
| reproductive shoot system mineral and ion content trait | 13 |
| cell or tissue mineral and ion content trait | 14 |
| **metabolite content trait** |  |
| carbohydrate content trait | 13 |
| lipid content trait | 1 |
| phenolic compound content trait | 5 |
| pigment content trait | 9 |
| protein content trait | 4 |
| **stable isotope ratio determination** | 14 |
| **plant morphology trait** |  |
| **whole plant morphology trait** | 5 |
| plant embryo morphology trait | 8 |
| **plant structure morphology trait** | 6 |
| portion of plant tissue morphology trait | 11 |
| plant cell morphology trait | 6 |
| **leaf morphology trait** | 30 |
| leaf size trait | 11 |
| leaf mass trait | 16 |
| leaf shape trait | 10 |
| leaf position trait | 5 |
| leaf stomatal complex morphology trait | 6 |
| leaf optical properties trait | 4 |
| **stem morphology trait** | 34 |
| stem mass trait | 6 |
| **bark morphology trait** | 9 |
| **root system morphology trait** | 14 |
| **reproductive shoot system morphology trait** | 3 |
| floral organ morphology trait | 4 |
| perianth morphology trait | 12 |
| androecium morphology trait | 14 |
| gynoecium morphology trait | 8 |
| fruit morphology trait | 13 |
| seed morphology trait | 16 |
| **vascular tissue morphology trait** | 5 |
| leaf vein morphology trait | 4 |
| xylem vessel morphology trait | 10 |
| **plant structure strength trait** | 10 |
| **biological process trait, physiological process trait** |  |
| photosynthetic trait | 2 |
| gas exchange trait | 13 |
| photosynthetic rate trait | 7 |
| respiration rate trait | 4 |
| transpiration rate trait | 6 |
| carbon dioxide concentration trait | 6 |
| photosystem performance trait | 11 |
| water transport trait, hydraulic trait | 44 |
| nutrient recycling trait | 2 |
| **life history trait** | 5 |
| whole plant phenotype trait | 16 |
| plant phenological trait | 16 |
| interspecific interactions trait | 8 |
| genetic structure trait | 2 |
| reproductive structure life history trait | 14 |
| fire response trait | 32 |
| chemical stress sensitivity trait | 3 |
| environmental tolerance trait | 16 |

\* The total number of traits in this table is greater than the total number of traits in APD, since some traits appear in multiple categories.

**Supplementary Table 2.**
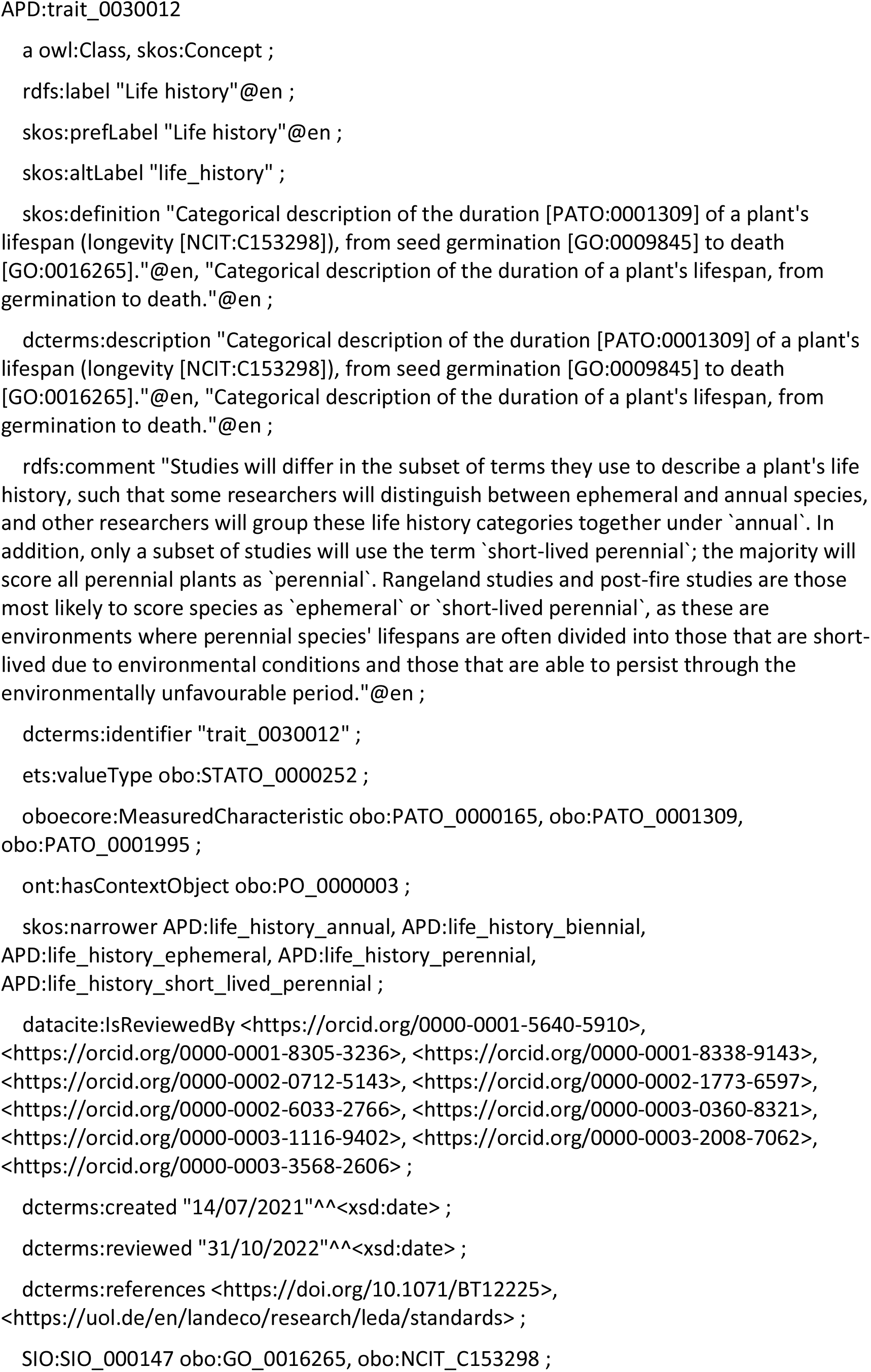
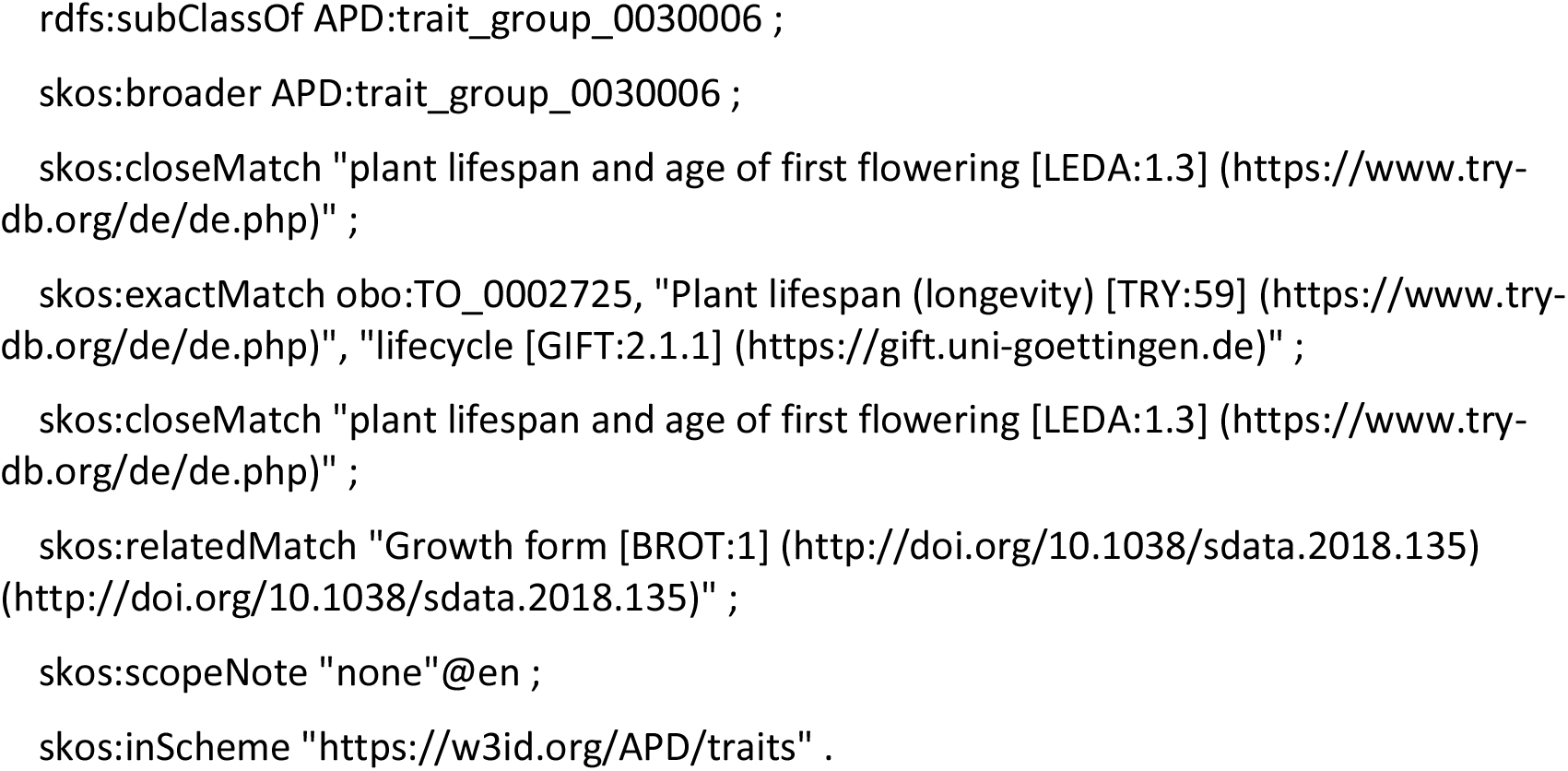
Output for the trait ‘life history’ from APD.ttl

**Supplementary Table 3.** Traits within the APD.

| label | alternate label (AusTraits `trait_name`) | APD identifier |
| --- | --- | --- |
| **Biochemical Traits** |  |  |
| Leaf aluminium (Al) content per unit leaf dry mass | leaf_Al_per_dry_mass | trait_0000012 |
| Leaf boron (B) content per unit leaf dry mass | leaf_B_per_dry_mass | trait_0000014 |
| Leaf carbon (C) content per unit leaf dry mass | leaf_C_per_dry_mass | trait_0000016 |
| Leaf calcium (Ca) content per unit leaf dry mass | leaf_Ca_per_dry_mass | trait_0000018 |
| Leaf chlorine (Cl) content per unit leaf dry mass | leaf_Cl_per_dry_mass | trait_0000020 |
| Leaf chromium (Cr) content per unit leaf dry mass | leaf_Cr_per_dry_mass | trait_0000022 |
| Leaf cobalt (Co) content per unit leaf dry mass | leaf_Co_per_dry_mass | trait_0000024 |
| Leaf copper (Cu) content per unit leaf dry mass | leaf_Cu_per_dry_mass | trait_0000026 |
| Leaf iron (Fe) content per unit leaf dry mass | leaf_Fe_per_dry_mass | trait_0000028 |
| Leaf potassium (K) content per unit leaf area | leaf_K_per_area | trait_0000029 |
| Leaf potassium (K) content per unit leaf dry mass | leaf_K_per_dry_mass | trait_0000030 |
| Leaf magnesium (Mg) content per unit leaf dry mass | leaf_Mg_per_dry_mass | trait_0000032 |
| Leaf manganese (Mn) content per unit leaf dry mass | leaf_Mn_per_dry_mass | trait_0000034 |
| Leaf molybdenum (Mo) content per unit leaf dry mass | leaf_Mo_per_dry_mass | trait_0000036 |
| Leaf nitrogen (N) content per unit leaf area | leaf_N_per_area | trait_0000037 |
| Leaf nitrogen (N) content per unit leaf dry mass | leaf_N_per_dry_mass | trait_0000038 |
| Leaf sodium (Na) content per unit leaf dry mass | leaf_Na_per_dry_mass | trait_0000040 |
| Leaf nickel (Ni) content per unit leaf dry mass | leaf_Ni_per_dry_mass | trait_0000042 |
| Leaf phosphorus (P) content per unit leaf area | leaf_P_per_area | trait_0000043 |
| Leaf phosphorus (P) content per unit leaf dry mass | leaf_P_per_dry_mass | trait_0000044 |
| Leaf sulphur (S) content per unit leaf dry mass | leaf_S_per_dry_mass | trait_0000046 |
| Leaf selenium (Se) content per unit leaf dry mass | leaf_Se_per_dry_mass | trait_0000048 |
| Leaf silicon (Si) content per unit leaf dry mass | leaf_Si_per_dry_mass | trait_0000050 |
| Leaf zinc (Zn) content per unit leaf dry mass | leaf_Zn_per_dry_mass | trait_0000052 |
| Leaf carbon to nitrogen ratio (C/N) | leaf_CN_ratio | trait_0000090 |
| Leaf nitrogen to phosphorus ratio (N/P) per unit leaf dry mass | leaf_NP_ratio | trait_0000091 |
| Senesced leaf aluminium (Al) content per unit leaf dry mass | leaf_senesced_Al_per_dry_mass | trait_0000112 |
| Senesced leaf boron (B) content per unit leaf dry mass | leaf_senesced_B_per_dry_mass | trait_0000114 |
| Senesced leaf carbon (C) content per unit leaf dry mass | leaf_senesced_C_per_dry_mass | trait_0000116 |
| Senesced leaf calcium (Ca) content per unit leaf dry mass | leaf_senesced_Ca_per_dry_mass | trait_0000118 |
| Senesced leaf copper (Cu) content per unit leaf dry mass | leaf_senesced_Cu_per_dry_mass | trait_0000126 |
| Senesced leaf iron (Fe) content per unit leaf dry mass | leaf_senesced_Fe_per_dry_mass | trait_0000128 |
| Senesced leaf potassium (K) content per unit leaf dry mass | leaf_senesced_K_per_dry_mass | trait_0000130 |
| Senesced leaf magnesium (Mg) content per unit leaf dry mass | leaf_senesced_Mg_per_dry_mass | trait_0000132 |
| Senesced leaf manganese (Mn) content per unit leaf dry mass | leaf_senesced_Mn_per_dry_mass | trait_0000134 |
| Senesced leaf molybdenum (Mo) content per unit leaf dry mass | leaf_senesced_Mo_per_dry_mass | trait_0000136 |
| Senesced leaf nitrogen (N) content per unit leaf dry mass | leaf_senesced_N_per_dry_mass | trait_0000138 |
| Senesced leaf sodium (Na) content per unit leaf dry mass | leaf_senesced_Na_per_dry_mass | trait_0000140 |
| Senesced leaf nickel (Ni) content per unit leaf dry mass | leaf_senesced_Ni_per_dry_mass | trait_0000142 |
| Senesced leaf phosphorus (P) content per unit leaf dry mass | leaf_senesced_P_per_dry_mass | trait_0000144 |
| Senesced leaf sulphur (S) content per unit leaf dry mass | leaf_senesced_S_per_dry_mass | trait_0000146 |
| Senesced leaf zinc (Zn) content per unit leaf dry mass | leaf_senesced_Zn_per_dry_mass | trait_0000152 |
| Leaf nitrogen resorption | leaf_N_resorption | trait_0022012 |
| Leaf phosphorus resorption | leaf_P_resorption | trait_0022013 |
| Stem carbon (C) content per unit stem dry mass | stem_C_per_dry_mass | trait_0000216 |
| Stem nitrogen (N) content per unit stem dry mass | stem_N_per_dry_mass | trait_0000238 |
| Wood carbon (C) content per unit wood dry mass | wood_C_per_dry_mass | trait_0000416 |
| Wood calcium (Ca) content per unit wood dry mass | wood_Ca_per_dry_mass | trait_0000418 |
| Wood potassium (K) content per unit wood dry mass | wood_K_per_dry_mass | trait_0000430 |
| Wood magnesium (Mg) content per unit wood dry mass | wood_Mg_per_dry_mass | trait_0000432 |
| Wood nitrogen (N) content per unit wood dry mass | wood_N_per_dry_mass | trait_0000438 |
| Wood sodium (Na) content per unit wood dry mass | wood_Na_per_dry_mass | trait_0000440 |
| Wood phosphorus (P) content per unit wood dry mass | wood_P_per_dry_mass | trait_0000444 |
| Dead wood calcium (Ca) content per unit dead wood dry mass | wood_dead_Ca_per_dry_mass | trait_0000518 |
| Dead wood potassium (K) content per unit dead wood dry mass | wood_dead_K_per_dry_mass | trait_0000530 |
| Dead wood magnesium (Mg) content per unit dead wood dry mass | wood_dead_Mg_per_dry_mass | trait_0000532 |
| Dead wood nitrogen (N) content per unit dead wood dry mass | wood_dead_N_per_dry_mass | trait_0000538 |
| Dead wood sodium (Na) content per unit dead wood dry mass | wood_dead_Na_per_dry_mass | trait_0000540 |
| Dead wood phosphorus (P) content per unit dead wood dry mass | wood_dead_P_per_dry_mass | trait_0000544 |
| Bark aluminium (Al) content per unit bark dry mass | bark_Al_per_dry_mass | trait_0000612 |
| Bark boron (B) content per unit bark dry mass | bark_B_per_dry_mass | trait_0000614 |
| Bark carbon (C) content per unit bark dry mass | bark_C_per_dry_mass | trait_0000616 |
| Bark calcium (Ca) content per unit bark dry mass | bark_Ca_per_dry_mass | trait_0000618 |
| Bark copper (Cu) content per unit bark dry mass | bark_Cu_per_dry_mass | trait_0000626 |
| Bark iron (Fe) content per unit bark dry mass | bark_Fe_per_dry_mass | trait_0000628 |
| Bark potassium (K) content per unit bark dry mass | bark_K_per_dry_mass | trait_0000630 |
| Bark magnesium (Mg) content per unit bark dry mass | bark_Mg_per_dry_mass | trait_0000632 |
| Bark manganese (Mn) content per unit bark dry mass | bark_Mn_per_dry_mass | trait_0000634 |
| Bark nitrogen (N) content per unit bark dry mass | bark_N_per_dry_mass | trait_0000638 |
| Bark sodium (Na) content per unit bark dry mass | bark_Na_per_dry_mass | trait_0000640 |
| Bark phosphorus (P) content per unit bark dry mass | bark_P_per_dry_mass | trait_0000644 |
| Bark sulphur (S) content per unit bark dry mass | bark_S_per_dry_mass | trait_0000646 |
| Bark zinc (Zn) content per unit bark dry mass | bark_Zn_per_dry_mass | trait_0000652 |
| Root carbon (C) content per unit root dry mass | root_C_per_dry_mass | trait_0000816 |
| Root nitrogen (N) content per unit root dry mass | root_N_per_dry_mass | trait_0000838 |
| Root phosphorus (P) content per unit root dry mass | root_P_per_dry_mass | trait_0000844 |
| Flower nitrogen (N) content per unit flower dry mass | flower_N_per_dry_mass | trait_0001038 |
| Fruit calcium (Ca) content per unit fruit dry mass | fruit_Ca_per_dry_mass | trait_0001118 |
| Fruit potassium (K) content per unit fruit dry mass | fruit_K_per_dry_mass | trait_0001130 |
| Fruit magnesium (Mg) content per unit fruit dry mass | fruit_Mg_per_dry_mass | trait_0001132 |
| Fruit nitrogen (N) content per unit fruit dry mass | fruit_N_per_dry_mass | trait_0001138 |
| Fruit phosphorus (P) content per unit fruit dry mass | fruit_P_per_dry_mass | trait_0001144 |
| Fruit sulphur (S) content per unit fruit dry mass | fruit_S_per_dry_mass | trait_0001146 |
| Seed calcium (Ca) content per unit seed dry mass | seed_Ca_per_seed_dry_mass | trait_0001218 |
| Seed potassium (K) content per unit seed dry mass | seed_K_per_seed_dry_mass | trait_0001230 |
| Seed magnesium (Mg) content per unit seed dry mass | seed_Mg_per_seed_dry_mass | trait_0001232 |
| Seed nitrogen (N) content per unit seed dry mass | seed_N_per_seed_dry_mass | trait_0001238 |
| Seed phosphorus (P) content per unit seed dry mass | seed_P_per_seed_dry_mass | trait_0001244 |
| Seed sulphur (S) content per unit seed dry mass | seed_S_per_seed_dry_mass | trait_0001246 |
| Leaf cell wall nitrogen (N) per unit cell wall dry mass | leaf_cell_wall_N_per_cell_wall_dry_mass | trait_0001511 |
| Leaf cell wall nitrogen (N) per unit leaf N content | leaf_cell_wall_N_per_leaf_N | trait_0001512 |
| Leaf rubisco nitrogen (N) content per unit leaf N content | leaf_rubisco_N_per_total_leaf_N | trait_0001513 |
| Leaf thylakoid protein nitrogen (N) content per unit leaf N content | leaf_thylakoid_N_per_total_leaf_N | trait_0001514 |
| Leaf epidermis calcium (Ca) content per unit leaf fresh mass | leaf_epidermis_Ca_per_fresh_mass | trait_0001611 |
| Leaf hypodermis calcium (Ca) content per unit leaf fresh mass | leaf_hypodermis_Ca_per_fresh_mass | trait_0001612 |
| Leaf internal parenchyma cell calcium (Ca) content per unit leaf fresh mass | leaf_internal_parenchyma_Ca_per_fresh_mass | trait_0001613 |
| Leaf palisade mesophyll cell calcium (Ca) content per unit leaf fresh mass | leaf_palisade_mesophyll_Ca_per_fresh_mass | trait_0001614 |
| Leaf sclerenchyma cell calcium (Ca) content per unit leaf fresh mass | leaf_sclerenchyma_Ca_per_fresh_mass | trait_0001615 |
| Leaf spongy mesophyll cell calcium (Ca) content per unit leaf fresh mass | leaf_spongy_mesophyll_Ca_per_fresh_mass | trait_0001616 |
| Leaf epidermis phosphorus (P) content per unit leaf fresh mass | leaf_epidermis_P_per_fresh_mass | trait_0001661 |
| Leaf hypodermis phosphorus (P) content per unit leaf fresh mass | leaf_hypodermis_P_per_fresh_mass | trait_0001662 |
| Leaf internal parenchyma cell phosphorus (P) content per unit leaf fresh mass | leaf_internal_parenchyma_P_per_fresh_mass | trait_0001663 |
| Leaf palisade mesophyll cell phosphorus (P) content per unit leaf fresh mass | leaf_palisade_mesophyll_P_per_fresh_mass | trait_0001664 |
| Leaf sclerenchyma cell phosphorus (P) content per unit leaf fresh mass | leaf_sclerenchyma_P_per_fresh_mass | trait_0001665 |
| Leaf spongy mesophyll cell phosphorus (P) content per unit leaf fresh mass | leaf_spongy_mesophyll_P_per_fresh_mass | trait_0001666 |
| Leaf total non-structural carbohydrate content per unit leaf area | leaf_total_non-structural_carbohydrates_per_area | trait_0002021 |
| Leaf total non-structural carbohydrate content per unit leaf dry mass | leaf_total_non-structural_carbohydrates_per_mass | trait_0002022 |
| Leaf cellulose content per unit leaf dry mass | leaf_cellulose_per_dry_mass | trait_0002024 |
| Leaf starch content per unit leaf area | leaf_starch_per_area | trait_0002025 |
| Leaf soluble starch content per unit leaf area | leaf_soluble_starch_per_area | trait_0002027 |
| Leaf soluble starch content per unit leaf dry mass | leaf_soluble_starch_per_mass | trait_0002028 |
| Leaf soluble sugar content per unit leaf area | leaf_soluble_sugars_per_area | trait_0002031 |
| Leaf soluble sugar content per unit leaf dry mass | leaf_soluble_sugars_per_mass | trait_0002032 |
| Leaf soluble protein content per unit leaf area | leaf_soluble_protein_per_area | trait_0002035 |
| Leaf insoluble protein content per unit leaf area | leaf_insoluble_protein_per_area | trait_0002037 |
| Leaf lignin content per unit leaf dry mass | leaf_lignin_per_dry_mass | trait_0002050 |
| Total leaf phenolic content per unit leaf dry mass | leaf_phenol_per_dry_mass | trait_0002052 |
| Leaf tannin content per unit leaf dry mass | leaf_tannin_per_dry_mass | trait_0002054 |
| Leaf carotenoid content per unit leaf area | leaf_carotenoid_per_area | trait_0002055 |
| Leaf carotenoid content per unit leaf dry mass | leaf_carotenoid_per_dry_mass | trait_0002056 |
| Leaf total chlorophyll content (chlorophyll A + B) per unit leaf area | leaf_chlorophyll_per_area | trait_0002081 |
| Leaf total chlorophyll content (chlorophyll A + B) per unit leaf dry mass | leaf_chlorophyll_per_dry_mass | trait_0002082 |
| Leaf chlorophyll A content per unit leaf area | leaf_chlorophyll_A_per_area | trait_0002083 |
| Leaf chlorophyll A content per unit leaf dry mass | leaf_chlorophyll_A_per_dry_mass | trait_0002084 |
| Leaf chlorophyll B content per unit leaf area | leaf_chlorophyll_B_per_area | trait_0002085 |
| Leaf chlorophyll B content per unit leaf dry mass | leaf_chlorophyll_B_per_dry_mass | trait_0002086 |
| Ratio of leaf chlorophyll A content to leaf chlorophyll B content | leaf_chlorophyll_A_B_ratio | trait_0002087 |
| Leaf rubisco content per unit leaf dry mass | leaf_rubisco_per_leaf_dry_mass | trait_0002090 |
| Stem soluble starch content per unit stem dry mass | stem_soluble_starch_per_mass | trait_0002127 |
| Stem soluble sugar content per unit stem dry mass | stem_soluble_sugars_per_mass | trait_0002131 |
| Bark cellulose content per unit bark dry mass | bark_cellulose_per_dry_mass | trait_0002224 |
| Bark lignin content per unit bark dry mass | bark_lignin_per_dry_mass | trait_0002250 |
| Bark tannin content per unit bark dry mass | bark_tannin_per_dry_mass | trait_0002255 |
| Root soluble starch content per unit root dry mass | root_soluble_starch_per_mass | trait_0002327 |
| Root soluble sugar content per unit root dry mass | root_soluble_sugars_per_mass | trait_0002331 |
| Seed protein content per unit seed dry mass | seed_protein_per_seed_dry_mass | trait_0002534 |
| Seed oil content per unit seed dry mass | seed_oil_per_seed_dry_mass | trait_0002544 |
| Leaf ash content per unit leaf dry mass | leaf_ash_per_dry_mass | trait_0002822 |
| Bark ash content per unit bark dry mass | bark_ash_per_dry_mass | trait_0002824 |
| Bark stable carbon isotope composition (delta13C) | bark_delta13C | trait_0003011 |
| Leaf stable carbon isotope composition (delta13C) | leaf_delta13C | trait_0003012 |
| Stem stable carbon isotope composition (delta13C) | stem_delta13C | trait_0003013 |
| Root stable carbon isotope composition (delta13C) | root_delta13C | trait_0003014 |
| Wood stable carbon isotope composition (delta13C) | wood_delta13C | trait_0003015 |
| Bark stable nitrogen isotope composition (delta15N) | bark_delta15N | trait_0003031 |
| Leaf stable nitrogen isotope composition (delta15N) | leaf_delta15N | trait_0003032 |
| Stem stable nitrogen isotope composition (delta15N) | stem_delta15N | trait_0003033 |
| Root stable nitrogen isotope composition (delta15N) | root_delta15N | trait_0003034 |
| Wood stable nitrogen isotope composition (delta15N) | wood_delta15N | trait_0003035 |
| Leaf xylem stable nitrogen isotope composition (delta15N) | leaf_xylem_delta15N | trait_0003052 |
| Root xylem stable nitrogen isotope composition (delta15N) | root_xylem_delta15N | trait_0003053 |
| Leaf stable oxygen isotope composition (delta18O) | leaf_delta18O | trait_0003072 |
| Stem water stable oxygen isotope composition (delta18O) | stem_water_delta18O | trait_0003092 |
| **Plant Morphology Trait** |  |  |
| Plant canopy width | plant_width | trait_0010021 |
| Plant canopy breadth | plant_breadth | trait_0010022 |
| Plant vegetative height | plant_height | trait_0010023 |
| Stem diameter at breast height | plant_diameter_breast_height | trait_0010024 |
| Stem count | stem_count | trait_0010025 |
| Plant spinescence | plant_spinescence | trait_0010070 |
| Embryo colour | embryo_colour | trait_0010110 |
| Cotyledon function | cotyledon_function | trait_0010111 |
| Cotyledon position at germination | cotyledon_position | trait_0010112 |
| Cotyledon hairiness | cotyledon_hairs | trait_0010113 |
| Hypocotyl hairiness | seedling_hypocotyl_hairs | trait_0010114 |
| Seedling first true leaf type | seedling_first_node_leaf_type | trait_0010160 |
| Seedling first node leaf count | seedling_first_node_leaf_count | trait_0010161 |
| Seedling germination location | seedling_germination_location | trait_0010162 |
| Leaf area | leaf_area | trait_0011211 |
| Leaflet area | leaflet_area | trait_0011212 |
| Leaf length | leaf_length | trait_0011213 |
| Leaf width | leaf_width | trait_0011214 |
| Leaf thickness | leaf_thickness | trait_0011215 |
| Leaf dry mass | leaf_dry_mass | trait_0011216 |
| Leaflet dry mass | leaflet_dry_mass | trait_0011217 |
| Leaf fresh mass | leaf_fresh_mass | trait_0011218 |
| Petiole length | petiole_length | trait_0011219 |
| Petiole width | petiole_width | trait_0011220 |
| Leaf mass per area | leaf_mass_per_area | trait_0011230 |
| Leaf lamina mass per area | leaf_lamina_mass_per_area | trait_0011231 |
| Leaf tissue density | leaf_density | trait_0011232 |
| Leaf area ratio (LAR) | leaf_area_ratio | trait_0011260 |
| Leaf mass fraction | leaf_mass_fraction | trait_0011261 |
| Leaf dry matter content (LDMC) | leaf_dry_matter_content | trait_0011262 |
| Leaf fresh mass per leaf area | leaf_fresh_mass_per_area | trait_0011263 |
| Leaf water content per unit leaf area (leaf succulence) | leaf_water_content_per_area | trait_0011264 |
| Leaf water content per unit leaf dry mass | leaf_water_content_per_dry_mass | trait_0011265 |
| Leaf water content per unit leaf fresh mass | leaf_water_content_per_fresh_mass | trait_0011266 |
| Leaf water content per unit saturated leaf mass | leaf_water_content_per_saturated_mass | trait_0011267 |
| Leaf cell wall fraction | leaf_cell_wall_fraction | trait_0011268 |
| Leaf type | leaf_type | trait_0011310 |
| Leaf shape | leaf_shape | trait_0011311 |
| Leaf base shape | leaf_base_shape | trait_0011312 |
| Leaf margin | leaf_margin | trait_0011313 |
| Leaf margin posture | leaf_margin_posture | trait_0011314 |
| Leaf lobation | leaf_lobation | trait_0011315 |
| Leaf compoundness | leaf_compoundness | trait_0011316 |
| Leaf divisions | leaf_lamina_division | trait_0011317 |
| Leaf lamina posture (leaf 3-dimensionality) | leaf_posture_numeric | trait_0011318 |
| Leaf lamina posture (leaf 3-dimensional shape) | leaf_lamina_posture | trait_0011319 |
| Leaf glaucousness | leaf_glaucousness | trait_0011360 |
| Mature leaf hairiness | leaf_hairs_adult_leaves | trait_0011361 |
| Juvenile phase leaf hairiness | leaf_hairs_juvenile_leaves | trait_0011362 |
| Immature leaf hairiness | leaf_hairs_immature_leaves | trait_0011363 |
| Leaf phyllotaxis | leaf_phyllotaxis | trait_0011410 |
| Leaf arrangement | leaf_arrangement | trait_0011411 |
| Leaf axil angle | leaf_axil_angle | trait_0011412 |
| Leaf inclination angle | leaf_inclination_angle | trait_0011413 |
| Leaf pendulousness | leaf_pendulousness | trait_0011414 |
| Cuticle thickness on the lower leaf surface | leaf_cuticle_thickness_abaxial | trait_0011510 |
| Cuticle thickness on the upper leaf surface | leaf_cuticle_thickness_adaxial | trait_0011511 |
| Leaf epidermis thickness | leaf_epidermis_thickness | trait_0011512 |
| Lower leaf side epidermis thickness | leaf_epidermis_thickness_abaxial | trait_0011513 |
| Upper leaf side epidermis thickness | leaf_epidermis_thickness_adaxial | trait_0011514 |
| Average leaf epidermal cell density | leaf_epidermal_cell_density_both_sides | trait_0011515 |
| Lower leaf side epidermal cell density | leaf_epidermal_cell_density_abaxial | trait_0011516 |
| Upper leaf side epidermal cell density | leaf_epidermal_cell_density_adaxial | trait_0011517 |
| Lower leaf side hypodermis thickness | leaf_hypodermis_thickness_abaxial | trait_0011518 |
| Upper leaf side hypodermis thickness | leaf_hypodermis_thickness_adaxial | trait_0011519 |
| Lower palisade mesophyll thickness | leaf_palisade_tissue_thickness_abaxial | trait_0011520 |
| Upper palisade mesophyll thickness | leaf_palisade_tissue_thickness_adaxial | trait_0011521 |
| Palisade cell length | leaf_palisade_cell_length | trait_0011522 |
| Palisade cell width | leaf_palisade_cell_width | trait_0011523 |
| Number of layers of palisade cells | leaf_palisade_layer_number | trait_0011524 |
| Spongy mesophyll cell thickness | leaf_spongy_mesophyll_thickness | trait_0011525 |
| Cell cross-sectional area | cell_cross-sectional_area | trait_0011526 |
| Stomatal density on the lower leaf surface | leaf_stomatal_density_abaxial | trait_0011610 |
| Stomatal density on the upper leaf surface | leaf_stomatal_density_adaxial | trait_0011611 |
| Stomatal density averaged across both leaf surfaces | leaf_stomatal_density_average | trait_0011612 |
| Stomatal distribution | leaf_stomatal_distribution | trait_0011613 |
| Stomatal hairiness | leaf_stomatal_hairs | trait_0011614 |
| Guard cell length | leaf_guard_cell_length | trait_0011615 |
| Leaf visible light transmission | leaf_transmission | trait_0011710 |
| Leaf visible light absorption | leaf_absorption | trait_0011711 |
| Leaf visible light reflection | leaf_reflectance | trait_0011712 |
| Leaf infra-red light reflection | leaf_reflectance_near_infrared | trait_0011713 |
| Stem cross-sectional area | stem_cross_sectional_area | trait_0011811 |
| Wood cross-sectional area | sapwood_cross_sectional_area | trait_0011812 |
| Terminal twig cross-sectional area | branch_terminal_twig_cross_sectional_area | trait_0011813 |
| Terminal twig length | branch_terminal_twig_length | trait_0011814 |
| Wood density | wood_density | trait_0011815 |
| Herbaceous stem density | stem_density | trait_0011816 |
| Huber value | huber_value | trait_0011911 |
| Leaf dry mass to stem dry mass ratio | leaf_mass_to_stem_mass_ratio | trait_0011912 |
| Stem dry mass to vegetative shoot dry mass ratio (support fraction) | stem_mass_to_shoot_mass_ratio | trait_0011913 |
| Side branch dry mass to whole plant dry mass ratio | branch_mass_fraction | trait_0011914 |
| Stem dry matter content (SDMC) | stem_dry_matter_content | trait_0011915 |
| Stem water content per unit saturated stem mass | stem_water_content_per_saturated_mass | trait_0011916 |
| Stem mass fraction | stem_mass_fraction | trait_0011917 |
| Bark morphology, Eucalyptus | bark_morphology_eucalyptus | trait_0012010 |
| Bark thickness | bark_thickness | trait_0012011 |
| Scaled bark thickness | bark_thickness_index | trait_0012012 |
| Bark density | bark_density | trait_0012013 |
| Bark dry mass per unit bark surface area | bark_dry_mass_per_surface_area | trait_0012014 |
| Bark water content per unit bark dry mass | bark_water_content_per_dry_mass | trait_0012015 |
| Bark water content per unit saturated bark mass | bark_water_content_per_saturated_mass | trait_0012016 |
| Root diameter | root_diameter | trait_0012111 |
| Root system morphology | root_system_classification | trait_0012112 |
| Fine root volume to coarse root volume ratio | root_fine_root_coarse_root_ratio | trait_0012113 |
| Root biomass depth distribution coefficient | root_distribution_coefficient | trait_0012114 |
| Root system type (presence of taproot) | root_system_type | trait_0012115 |
| Specific root length (SRL) | root_specific_root_length | trait_0012116 |
| Specific tap root length (STRL) | root_specific_taproot_length | trait_0012117 |
| Root surface area per unit root dry mass (specific root area) | root_specific_root_area | trait_0012118 |
| Root wood density | root_wood_density | trait_0012119 |
| Root to shoot ratio | root_shoot_ratio | trait_0012120 |
| Root dry matter content (RDMC) | root_dry_matter_content | trait_0012121 |
| Root mass fraction | root_mass_fraction | trait_0012122 |
| Seed accessory cost fraction | accessory_cost_fraction | trait_0012221 |
| Seed accessory cost mass | accessory_cost_mass | trait_0012222 |
| Number of androecium parts in each whorl (Androecium structural merism) | flower_androecium_structural_merism | trait_0012410 |
| Androecium structural phyllotaxis | flower_androecium_structural_phyllotaxis | trait_0012411 |
| Number of androecium structural whorls | flower_androecium_structural_whorls_count | trait_0012412 |
| Anther attachment | flower_anther_attachment | trait_0012413 |
| Connective extension (apical) | flower_anther_connective_extension | trait_0012414 |
| Anther dehiscence | flower_anther_dehiscence | trait_0012415 |
| Anther orientation | flower_anther_orientation | trait_0012416 |
| Flower colour | flower_colour | trait_0012417 |
| Perianth colour | perianth_colour | trait_0012418 |
| Flower length | flower_length | trait_0012419 |
| Flower diameter | flower_diameter | trait_0012420 |
| Floral orientation | flower_orientation | trait_0012421 |
| Maximum flower number | flower_count_maximum | trait_0012422 |
| Number of fertile stamens | flower_fertile_stamens_count | trait_0012431 |
| Filament presence and shape | flower_filament | trait_0012432 |
| Fusion of filaments | flower_filament_fusion | trait_0012433 |
| Fusion of filaments to inner perianth series | flower_filament_fusion_to_inner_perianth | trait_0012434 |
| Gynoecium phyllotaxis | flower_gynoecium_phyllotaxis | trait_0012441 |
| Placentation | flower_gynoecium_placentation | trait_0012442 |
| Fusion of ovaries | flower_ovary_fusion | trait_0012443 |
| Ovary position | flower_ovary_position | trait_0012444 |
| Number of ovules per functional carpel | flower_ovules_per_functional_carpel_count | trait_0012445 |
| Perianth differentiation | flower_perianth_differentiation | trait_0012461 |
| Fusion of perianth | flower_perianth_fusion | trait_0012462 |
| Number of perianth parts in each whorl (Perianth merism) | flower_perianth_merism | trait_0012463 |
| Number of perianth parts | flower_perianth_parts_count | trait_0012464 |
| Perianth phyllotaxis | flower_perianth_phyllotaxis | trait_0012465 |
| Symmetry of perianth | flower_perianth_symmetry | trait_0012466 |
| Number of perianth whorls | flower_perianth_whorls_count | trait_0012467 |
| Pollen grain aperture shape | flower_pollen_aperture_shape | trait_0012471 |
| Number of pollen grain apertures | flower_pollen_apertures_count | trait_0012472 |
| Pollen grain length | flower_pollen_length | trait_0012473 |
| Number of structural carpels | flower_structural_carpels_count | trait_0012481 |
| Floral structural sex | flower_structural_sex_type | trait_0012482 |
| Style differentiation | flower_style_differentiation | trait_0012483 |
| Fusion of styles | flower_style_fusion | trait_0012484 |
| Fruit dry mass | fruit_dry_mass | trait_0012511 |
| Fruit length | fruit_length | trait_0012512 |
| Fruit width | fruit_width | trait_0012513 |
| Fruit breadth | fruit_height | trait_0012514 |
| Fruit wall thickness | fruit_wall_thickness | trait_0012515 |
| Fruit type | fruit_type | trait_0012516 |
| Fruit fleshiness | fruit_fleshiness | trait_0012517 |
| Fruit dehiscence | fruit_dehiscence | trait_0012518 |
| Fruit colour | fruit_colour | trait_0012519 |
| Seed dry mass | seed_dry_mass | trait_0012610 |
| Diaspore dry mass | diaspore_dry_mass | trait_0012611 |
| Seed embryo and endosperm dry mass | seed_dry_mass_reserve | trait_0012612 |
| Seed length | seed_length | trait_0012613 |
| Seed width | seed_width | trait_0012614 |
| Seed height | seed_height | trait_0012615 |
| Seed volume | seed_volume | trait_0012616 |
| Seed count | seed_count | trait_0012617 |
| Seed shape | seed_shape | trait_0012618 |
| Seed surface hairs | seed_surface_hairs | trait_0012619 |
| Seed surface texture | seed_surface_texture | trait_0012620 |
| Seed surface reflectivity | seed_surface_reflectivity | trait_0012621 |
| Diaspore fleshiness | diaspore_fleshiness | trait_0012622 |
| Dispersal appendage | dispersal_appendage | trait_0012623 |
| Dispersal unit | dispersal_unit | trait_0012624 |
| Leaf secondary vein angle | leaf_secondary_vein_angle | trait_0013011 |
| Major leaf vein density | leaf_major_vein_density | trait_0013012 |
| Length of all minor and major leaf lamina veins per unit area | leaf_total_vein_density | trait_0013013 |
| Leaf vein frequency | leaf_vein_frequency | trait_0013014 |
| Stem xylem vessel density | stem_vessel_density | trait_0013111 |
| Leaf xylem vessel density | leaf_vessel_density | trait_0013112 |
| Stem xylem vessel diameter | stem_vessel_diameter | trait_0013113 |
| Stem xylem vessel hydraulic mean diameter | stem_vessel_diameter_hydraulic | trait_0013114 |
| Leaf xylem vessel diameter | leaf_vessel_diameter | trait_0013115 |
| Stem xylem vessel lumen fraction | stem_vessel_lumen_fraction | trait_0013116 |
| Stem xylem vessel multiple fraction | stem_vessel_multiple_fraction | trait_0013117 |
| Stem non-lumen fraction | stem_vessel_non_lumen_fraction | trait_0013118 |
| xylem vessel wall fraction | stem_vessel_wall_fraction | trait_0013119 |
| Xylem vulnerability index | stem_xylem_vulnerability_index | trait_0013120 |
| Wood axial parenchyma fraction | wood_axial_parenchyma_fraction | trait_0013161 |
| Wood conduit fraction | wood_conduit_fraction | trait_0013162 |
| Wood fibre fraction | wood_fibre_fraction | trait_0013163 |
| Wood ray parenchyma fraction | wood_ray_parenchyma_fraction | trait_0013164 |
| Wood tracheid fraction | wood_tracheid_fraction | trait_0013165 |
| Leaf work to punch | leaf_work_to_punch | trait_0014011 |
| Leaf specific work to punch | leaf_work_to_punch_adjusted | trait_0014012 |
| Leaf work to shear | leaf_work_to_shear | trait_0014013 |
| Leaf specific work to shear (fracture toughness) | leaf_work_to_shear_adjusted | trait_0014014 |
| Leaf work to tear | leaf_work_to_tear | trait_0014015 |
| Leaf specific work to tear | leaf_work_to_tear_adjusted | trait_0014016 |
| Bark modulus of elasticity | bark_modulus_of_elasticity | trait_0014017 |
| Stem modulus of elasticity | stem_modulus_of_elasticity | trait_0014018 |
| Xylem modulus of elasticity | xylem_modulus_of_elasticity | trait_0014019 |
| Modulus of rupture | modulus_of_rupture | trait_0014020 |
| **Biological Process Trait (Physiological Process Trait)** |  |  |
| Plant photosynthetic pathway | photosynthetic_pathway | trait_0020221 |
| Bark photosynthesis | bark_photosynthetic_status | trait_0020222 |
| Leaf photosynthesis rate per unit leaf area under ambient light and CO2 (A) | leaf_photosynthetic_rate_per_area_ambient | trait_0020240 |
| Leaf photosynthesis rate per unit leaf area under saturating light and CO2 (Amax) | leaf_photosynthetic_rate_per_area_maximum | trait_0020241 |
| Leaf photosynthesis rate per unit leaf area under saturating light and ambient CO2 (Asat) | leaf_photosynthetic_rate_per_area_saturated | trait_0020242 |
| Leaf photosynthesis rate per unit leaf dry mass under ambient light and CO2 (A) | leaf_photosynthetic_rate_per_dry_mass_ambient | trait_0020243 |
| Leaf photosynthesis rate per unit leaf dry mass under saturating light and CO2 (Amax) | leaf_photosynthetic_rate_per_dry_mass_maximum | trait_0020244 |
| Leaf photosynthesis rate per unit leaf dry mass under saturating light and ambient CO2 (Asat) | leaf_photosynthetic_rate_per_dry_mass_saturated | trait_0020245 |
| Leaf internal CO2 concentration during Amax measurement (ci) | leaf_intercellular_CO2_concentration_at_Amax | trait_0020310 |
| Internal CO2 concentration during Asat measurement (ci) | leaf_intercellular_CO2_concentration_at_Asat | trait_0020311 |
| Internal CO2 concentration under ambient conditions (ci) | leaf_intercellular_CO2_concentration_at_Aambient | trait_0020312 |
| Ratio of internal to external CO2 concentrations (ci/ca) | leaf_intercellular_CO2_concentration_to_atmospheric_CO2_concentration_ratio | trait_0020313 |
| CO2 concentration inside chloroplasts (cc) | leaf_chloroplast_CO2_concentration | trait_0020314 |
| Ambient CO2 concentration (ca) | atmospheric_CO2_concentration | trait_0020315 |
| Leaf Jmax per unit leaf area (Jmax) | leaf_photosynthesis_Jmax_per_area | trait_0020410 |
| Leaf Jmax per unit leaf area at 25 deg C (Jmax25) | leaf_photosynthesis_Jmax_per_area_25C | trait_0020411 |
| Leaf Jmax per unit leaf mass (Jmax) | leaf_photosynthesis_Jmax_per_mass | trait_0020412 |
| Leaf Vcmax per unit leaf area (Vcmax) | leaf_photosynthesis_Vcmax_per_area | trait_0020413 |
| Leaf Vcmax per unit leaf area at 25 deg C (Vcmax25) | leaf_photosynthesis_Vcmax_per_area_25C | trait_0020414 |
| Leaf Vcmax per unit leaf mass (Vcmax) | leaf_photosynthesis_Vcmax_per_mass | trait_0020415 |
| Leaf Jmax to leaf Vcmax ratio at 25 deg C | leaf_photosynthesis_Jmax_over_Vcmax_25C | trait_0020416 |
| Leaf maximum quantum yield (Fv/Fm) | leaf_fluorescence_fv_over_fm | trait_0020417 |
| Leaf ambient quantum yield | leaf_fluorescence_quantum_yield | trait_0020418 |
| Leaf quantum yield, gas exchange measurement | leaf_gas_exchange_quantum_yield | trait_0020419 |
| Leaf respiration rate per unit leaf area, in the dark (Rdark) | leaf_dark_respiration_per_area | trait_0020510 |
| Leaf respiration rate per unit leaf dry mass, in the dark (Rdark) | leaf_dark_respiration_per_dry_mass | trait_0020511 |
| Leaf respiration rate per unit leaf area, in the light (Rday) | leaf_light_respiration_per_area | trait_0020512 |
| Stem respiration rate per unit stem area, in the dark | stem_dark_respiration_per_area | trait_0020513 |
| Leaf stomatal conductance to water vapour per unit leaf area under ambient conditions (gsw) | leaf_stomatal_conductance_per_area_ambient | trait_0020610 |
| Leaf stomatal conductance to water vapour per unit leaf area during Amax measurement (gsw) | leaf_stomatal_conductance_per_area_at_Amax | trait_0020611 |
| Leaf stomatal conductance to water vapour per unit leaf area during Asat measurement (gsw) | leaf_stomatal_conductance_per_area_at_Asat | trait_0020612 |
| Leaf stomatal water vapour resistance under ambient conditions | leaf_stomatal_resistance_ambient | trait_0020630 |
| Leaf mesophyll conductance to carbon dioxide per unit leaf area (gm) | leaf_mesophyll_conductance_per_area | trait_0020640 |
| Leaf mesophyll conductance to carbon dioxide per unit leaf mass (gm) | leaf_mesophyll_conductance_per_mass | trait_0020641 |
| Leaf transpiration per unit leaf area under ambient conditions (E) | leaf_transpiration_per_area_ambient | trait_0020660 |
| Leaf transpiration per unit leaf area during Amax measurement (E) | leaf_transpiration_per_area_at_Amax | trait_0020661 |
| Leaf transpiration per unit leaf area during Asat measurement (E) | leaf_transpiration_per_area_at_Asat | trait_0020662 |
| Leaf transpiration rate per unit leaf area, in the dark | leaf_dark_transpiration_per_area | trait_0020663 |
| Integrated plant transpiration | integrated_plant_transpiration | trait_0020664 |
| Whole plant sapflow | whole_plant_sapflow | trait_0020665 |
| Leaf photosynthetic nitrogen use efficiency during Amax measurement (PNUE) | leaf_photosynthetic_nitrogen_use_efficiency_maximum | trait_0020710 |
| Leaf photosynthetic nitrogen use efficiency during Asat measurement (PNUE) | leaf_photosynthetic_nitrogen_use_efficiency_saturated | trait_0020711 |
| Leaf photosynthetic phosphorus use efficiency during Amax measurement PPUE) | leaf_photosynthetic_phosphorus_use_efficiency_maximum | trait_0020712 |
| Leaf photosynthetic phosphorus use efficiency during Asat measurement (PPUE) | leaf_photosynthetic_phosphorus_use_efficiency_saturated | trait_0020713 |
| Integrated water use efficiency | leaf_water_use_efficiency_integrated | trait_0020760 |
| Intrinsic water use efficiency (WUEi) | leaf_water_use_efficiency_intrinsic | trait_0020761 |
| Instantaneous water use efficiency (WUE) | leaf_water_use_efficiency_instantaneous | trait_0020762 |
| Stem hydraulic conductivity (Kh) | stem_hydraulic_conductivity | trait_0021013 |
| Sapwood specific hydraulic conductivity (Ks) | sapwood_specific_hydraulic_conductivity | trait_0021014 |
| Theoretical sapwood specific hydraulic conductivity (Ks) | sapwood_specific_hydraulic_conductivity_theoretical | trait_0021015 |
| Stem specific hydraulic conductivity (Ks) | stem_specific_hydraulic_conductivity | trait_0021016 |
| Leaf specific hydraulic conductance (kleaf) | leaf_specific_hydraulic_conductance | trait_0021017 |
| Leaf specific hydraulic conductivity (Kl) | leaf_specific_hydraulic_conductivity | trait_0021018 |
| Root hydraulic conductivity (Kh) | root_hydraulic_conductivity | trait_0021021 |
| Root specific hydraulic conductivity | root_specific_hydraulic_conductivity | trait_0021022 |
| Pre-dawn water potential | water_potential_predawn | trait_0021030 |
| Midday water potential | water_potential_midday | trait_0021031 |
| Stem sapwood capacitance (C) | stem_sapwood_capacitance | trait_0021032 |
| Leaf capacitance (Cleaf) | leaf_capacitance | trait_0021033 |
| Root sapwood capacitance (C) | root_sapwood_capacitance | trait_0021034 |
| Leaf xylem pressure, 50% lost conductance | leaf_hydraulic_vulnerability | trait_0021050 |
| Stem xylem pressure, 12% lost conductivity | water_potential_12percent_lost_conductivity | trait_0021051 |
| Stem xylem pressure, 50% lost conductivity | water_potential_50percent_lost_conductivity | trait_0021052 |
| Stem xylem pressure, 88% lost conductivity | water_potential_88percent_lost_conductivity | trait_0021053 |
| Hydraulic safety margin, 50% | hydraulic_safety_margin_50 | trait_0021054 |
| Hydraulic safety margin, 88% | hydraulic_safety_margin_88 | trait_0021055 |
| Leaf turgor loss point | leaf_turgor_loss_point | trait_0021056 |
| Osmotic potential | osmotic_potential | trait_0021057 |
| Osmotic potential at full turgor | osmotic_potential_at_full_turgor | trait_0021058 |
| Bulk modulus of elasticity (e) | bulk_modulus_of_elasticity | trait_0021059 |
| Leaf relative water content predawn | leaf_relative_water_content_predawn | trait_0021060 |
| Leaf relative water content at turgor loss point | leaf_relative_water_content_at_turgor_loss_point | trait_0021061 |
| Root xylem pressure, 50% lost conductivity | root_water_potential_50percent_lost_conductivity | trait_0021062 |
| Photochemical reflectance index (PRI) | leaf_photochemical_reflectance_index | trait_0020813 |
| Water band index | leaf_water_band_index | trait_0020814 |
| Modified normalized difference vegetation index (modified NDVI) | modified_NDVI | trait_0020815 |
| Modified chlorophyll absorption ratio index 705 | leaf_chlorophyll_index_modified_ND705 | trait_0020816 |
| **Life History Trait** |  |  |
| Plant growth form | plant_growth_form | trait_0030010 |
| Life form | life_form | trait_0030011 |
| Life history | life_history | trait_0030012 |
| Ephemeral life history class | life_history_ephemeral_class | trait_0030013 |
| Lifespan | lifespan | trait_0030014 |
| Plant growth substrate | plant_growth_substrate | trait_0030015 |
| Plant photosynthetic organ | plant_photosynthetic_organ | trait_0030016 |
| Plant alternative energy and nutrient acquisition strategies | plant_alternative_energy_and_nutrient_acquisition_strategy | trait_0030017 |
| Woodiness | woodiness | trait_0030018 |
| Detailed woodiness categories | woodiness_detailed | trait_0030019 |
| Physical defence structures | plant_physical_defence_structures | trait_0030020 |
| Plant climbing mechanisms | plant_climbing_mechanism | trait_0030021 |
| Plant succulence | plant_succulence | trait_0030022 |
| Stem growth habit | stem_growth_habit | trait_0030023 |
| Leaf phenology | leaf_phenology | trait_0030024 |
| Leaf lifespan | leaf_lifespan | trait_0030025 |
| Competitive stratum | competitive_stratum | trait_0030026 |
| Plant nitrogen fixation capacity | nitrogen_fixing | trait_0030027 |
| Plant root structures | root_structure | trait_0030028 |
| Plant parasitism status | parasitic | trait_0030029 |
| Plant sex type | sex_type | trait_0030060 |
| Pollination syndrome | pollination_syndrome | trait_0030061 |
| Pollination system | pollination_system | trait_0030062 |
| Plant genome size | genome_size | trait_0030080 |
| Chromosome ploidy | ploidy | trait_0030081 |
| Age of reproductive maturity | reproductive_maturity | trait_0030210 |
| Diaspore dispersal syndrome | dispersal_syndrome | trait_0030211 |
| Diaspore dispersal agents | dispersers | trait_0030212 |
| Environmental flowering cues | flowering_cues | trait_0030213 |
| Flowering time, by month | flowering_time | trait_0030214 |
| Fruiting time, by month | fruiting_time | trait_0030215 |
| Seedling recruitment time, by month | recruitment_time | trait_0030216 |
| Seedling establishment conditions | seedling_establishment_conditions | trait_0030217 |
| Canopy light environment required for reproduction | reproductive_light_environment_index | trait_0030218 |
| Canopy light environment required for seedling establishment | establishment_light_environment_index | trait_0030219 |
| Seed storage location | seedbank_location | trait_0030411 |
| Serotiny | serotiny | trait_0030412 |
| Seedbank longevity class | seedbank_longevity_class | trait_0030413 |
| Seedbank longevity | seedbank_longevity | trait_0030414 |
| Dormancy type | seed_dormancy_class | trait_0030415 |
| Seed germination treatment | seed_germination_treatment | trait_0030416 |
| Seed germination proportion | seed_germination | trait_0030417 |
| Seed viability | seed_viability | trait_0030418 |
| Seed germination time | seed_germination_time | trait_0030419 |
| Vegetative reproduction ability | vegetative_reproduction_ability | trait_0030510 |
| Clonal spread mechanism | clonal_spread_mechanism | trait_0030511 |
| Storage organ | storage_organ | trait_0030512 |
| Bud bank location | bud_bank_location | trait_0030513 |
| Sprout depth | sprout_depth | trait_0030514 |
| Post-fire resprouting capacity | resprouting_capacity | trait_0030610 |
| Post-fire proportion resprouting individuals | resprouting_capacity_proportion_individuals | trait_0030611 |
| Post-fire resprouting capacity of juvenile plants | resprouting_capacity_juvenile | trait_0030612 |
| Time from seedling germination until individuals survive a fire | resprouting_capacity_time_from_germination | trait_0030613 |
| Post-fire to pre-fire stem ratio | resprouting_capacity_stem_ratio | trait_0030614 |
| Plant vegetative response to disturbances other than fire | resprouting_capacity_non_fire_disturbance | trait_0030615 |
| Fire exposure level | fire_exposure_level | trait_0030651 |
| Post-fire recruitment | post_fire_recruitment | trait_0030652 |
| Post-fire flowering | post_fire_flowering | trait_0030653 |
| Time from fire to first flowering | fire_time_from_fire_to_flowering | trait_0030654 |
| Time from fire until 50% of individuals are flowering | fire_time_from_fire_to_50_percent_flowering | trait_0030655 |
| Time from fire to peak flowering | fire_time_from_fire_to_peak_flowering | trait_0030656 |
| Time from fire until flowering declines | fire_time_from_fire_to_flowering_decline | trait_0030657 |
| Time from fire to fruiting | fire_time_from_fire_to_fruiting | trait_0030658 |
| Time from fire until 50% of individuals are fruiting | fire_time_from_fire_to_50_percent_fruiting | trait_0030659 |
| Fuel bed bulk density | fire_fuel_bed_bulk_density | trait_0030710 |
| Fuel consumption by fire | fire_fuel_consumption | trait_0030711 |
| Fire rate of spread | fire_rate_of_spread | trait_0030712 |
| Leaf smoulder duration | fire_smoulder_duration | trait_0030713 |
| Leaf flame duration | fire_flame_duration | trait_0030714 |
| Leaf flame and smoulder duration | fire_total_burn_duration | trait_0030715 |
| Fire time to ignition | fire_time_to_ignition | trait_0030716 |
| Plant resource requirements and tolerance | plant_type_by_resource_use | trait_0030810 |
| Plant flood regime response | plant_flood_regime_classification | trait_0030811 |
| Plant water-logging tolerance | plant_tolerance_water_logged_soils | trait_0030812 |
| Plant inundation tolerance | plant_tolerance_inundation | trait_0030813 |
| Plant snow tolerance | plant_tolerance_snow | trait_0030814 |
| Plant soil salinity tolerance | plant_tolerance_soil_salinity | trait_0030815 |
| Plant salt tolerance strategy | plant_tolerance_salt | trait_0030816 |
| Plant calcium sensitivity | plant_tolerance_calcicole | trait_0030817 |
| Plant fire tolerance strategy | plant_tolerance_fire | trait_0030818 |

**Supplementary Table 4.**
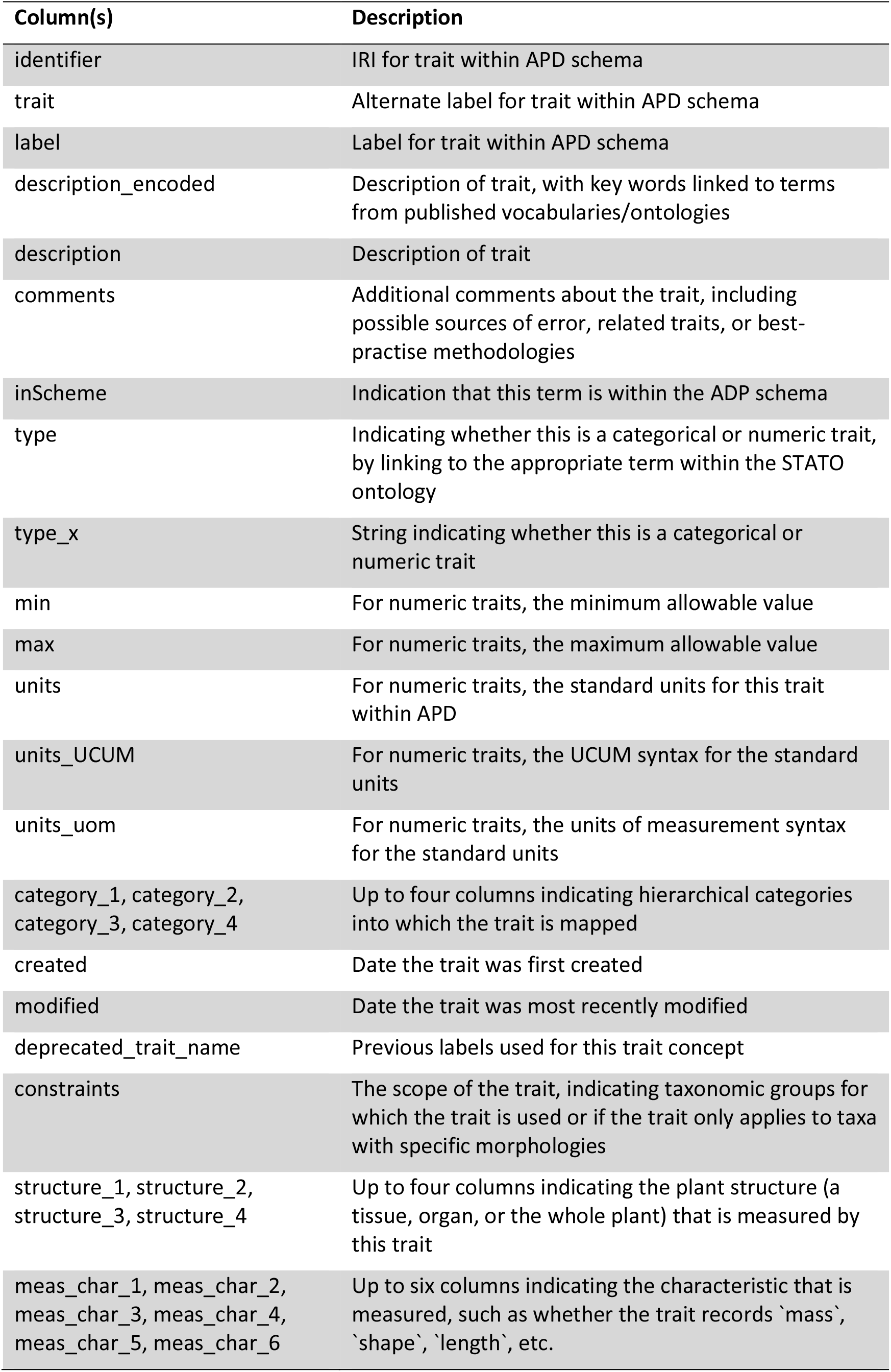
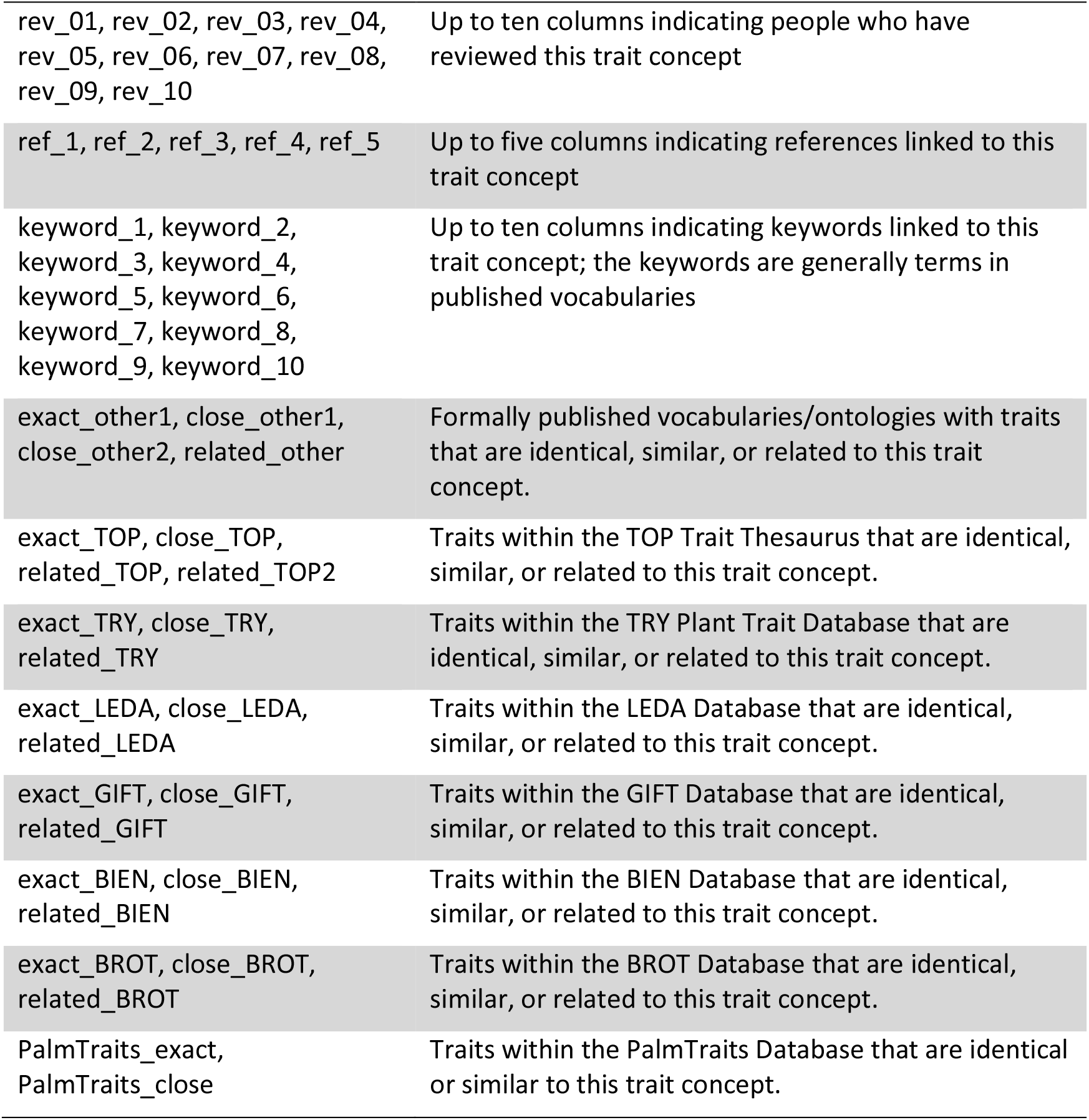
Columns in the data table APD_traits.csv

**Supplementary Table 5.** Columns in the data table APD_references.csv

| Column | Description | Annotation Property* |
| --- | --- | --- |
| Entity | URL for reference, if available |  |
| label | Author-year label for reference | skos:label |
| citation | Full reference citation | dcterms:bibliographicCitation |
| identifier | Identifier for reference, a DOI when available, or otherwise an ISBN (for books) or URL (for websites) | dcterms:identifier |
| title | The title of the reference | dcterms:title |

\* See Table 3 footnotes for the full schema URL's associated with each annotation property prefix.

**Supplementary Table 6.** Columns in the data table APD_reviewers.csv

| Column | Description | Annotation Property* |
| --- | --- | --- |
| Entity | URL for the reviewer’s ORCID profile |  |
| label | Reviewer’s full name | skos:label |
| ORCID | Reviewer’s ORCID number | obo:IAO_0000708 (ORCID identifier) |

\* See Table 3 footnotes for the full schema URL’s associated with each annotation property prefix.

**Supplementary Table 7.** Columns in the data table APD_units.csv

| Column | Description | Annotation Property* |
| --- | --- | --- |
| Entity | Units of Measurement IRI for the specific units of measurement |  |
| label | Label for these units of measurement | skos:label |
| altLabel | Alternative written label for these units of measurement | skos:altLabel |
| description | Verbal description of these units of measurement | dcterms:description |
| SI_code | The International System of Units code for these units of measurement | uom:SI_code |
| UCUM_code | The Unified Code for Units of Measure code for these units of measurement | uom:UCUM_code |
| exactMatch | Up to 6 columns indicating exact matches for this unit of measurement in other ontologies | skos:exactMatch |

\* See Table 3 footnotes for the full schema URL's associated with each annotation property prefix.

**Supplementary Table 8.** Columns in the data table published_classes.csv

| Column | Description | Annotation Property* |
| --- | --- | --- |
| Entity | URI for a specific term (class) in a published ontology |  |
| label | Label for the term | skos:label |
| description | Verbal description of the term | dcterms:description |
| identifier | Identifier for the term within a specific vocabulary | dcterms:identifier |
| inScheme | URI for the vocabulary in which the term is published | skos:inScheme |
| prefix | Prefix for the specific vocabulary | (Not used) |
| vocabulary | Name of the specific vocabulary | (Not used) |

\* See Table 3 footnotes for the full schema URL's associated with each annotation property prefix.

**Supplementary Table 9.** Columns in the data table APD_categorical_values.csv

| Column | Description | Annotation Property* |
| --- | --- | --- |
| identifier | Identifier for a specific categorical trait value within APD | dcterms:identifier |
| label | Label for the categorical trait value | skos:label |
| description | Description of the categorical trait value | dcterms:description |
| trait_name | Trait name to which the categorical trait value refers | skos:broader |

\* See Table 3 footnotes for the full schema URL's associated with each annotation property prefix.

**Supplementary Table 10.** Columns in the data table APD_trait_hierarchy.csv

| Column | Description | Annotation Property* |
| --- | --- | --- |
| Entity | URI for a trait category (hierarchical level) within APD |  |
| label | Label for a trait category (hierarchical level) within APD | skos:label |
| description | Description of a trait category (hierarchical level) within APD | dcterms:description |
| Parent | Superclass (higher hierarchical level) for a trait category within APD | skos:broader |
| exactMatch | Link to identical concept in a published ontology | skos:exactMatch |
| tier_1 | Highest hierarchical level into which the category fits | (Not used) |
| tier_2 | Second highest hierarchical level into which the category fits | (Not used) |
| tier_3 | Third highest hierarchical level into which the category fits (if applicable) | (Not used) |
| tier_4 | Fourth highest hierarchical level into which the category fits (if applicable) | (Not used) |
| hierarchy | Written string indicating the full hierarchy of the specific category | (Not used) |

\* See Table 3 footnotes for the full schema URL's associated with each annotation property prefix.

**Supplementary Table 11.** Columns in the data table APD_glossary.csv

| Column | Description | Annotation Property* |
| --- | --- | --- |
| identifier | Identifier for a specific glossary term within APD/glossary | dcterms:identifier |
| label | Label for the glossary term | skos:label |
| description | Description of the glossary term | dcterms:description |

\* See Table 3 footnotes for the full schema URL's associated with each annotation property prefix.

**Supplementary Table 12.** Columns in the data table APD_annotation_properties.csv

| Column | Description | Annotation Property* |
| --- | --- | --- |
| Entity | URI for annotation properties used by the APD |  |
| label | Label for annotation properties used by the APD, from its own vocabulary | skos:label |
| description | Description of annotation properties used by the APD, from its own vocabulary | dcterms:description |
| issued | Date a term was issued within its vocabulary | dcterms:issued |
| comment | Additional comments about annotation properties used by the APD, from its own vocabulary | rdfs:comment |
| isDefinedBy | URI for the vocabulary in which the term is published | rdfs:isDefinedBy |
| inScheme | URI for the vocabulary in which the term is published | skos:inScheme |

\* See Table 3 footnotes for the full schema URL's associated with each annotation property prefix.

**Supplementary Table 13.** Columns in the data table APD_namespace_declaration.csv

| Column | Description | Annotation Property* |
| --- | --- | --- |
| prefix | Prefix used for a specific vocabulary within APD machine-readable representations |  |
| Scheme | URI for each vocabulary used in the APD | skos:inScheme |

\* See Table 3 footnotes for the full schema URL's associated with each annotation property prefix.

**Supplementary Table 14.** Columns in the data table APD_resource.csv

| Column | Description |
| --- | --- |
| Subject | Entity URI for the APD schema |
| Predicate | Annotation properties for the APD schema |
| Object | Value of a particular annotation property for the APD schema |

## Notes

https://doi.org/10.5281/zenodo.8040789

https://vocabs.ardc.edu.au/viewById/649

https://w3id.org/APD

### Competing Interest Statement

The authors have declared no competing interest.

## References

1. Scheiter, S., Langan, L. & Higgins, S. I. Next-generation dynamic global vegetation models: learning from community ecology. New Phytol. 198, 957–969 (2013).

2. Gallagher, R. V. et al. A guide to using species trait data in conservation. One Earth 4, 927–936 (2021).

3. Garnier, E., Navas, M.-L. & Grigulis, K. Trait-based ecology: definitions, methods, and a conceptual framework. in Plant Functional Diversity: Organism traits, community structure, and ecosystem properties (eds. Garnier, E., Navas, M.-L. & Grigulis, K.) 0 (Oxford University Press, 2015). doi:10.1093/acprof:oso/9780198757368.003.0002.

4. De Queiroz, K. Species Concepts and Species Delimitation. Syst. Biol. 56, 879–886 (2007).

5. Gallagher, R. V. et al. Open Science principles for accelerating trait-based science across the Tree of Life. *Nat*. Ecol. Evol. 4, 294–303 (2020).

6. Kattge, J. et al. TRY plant trait database – enhanced coverage and open access. Glob. Change Biol. 26, 119–188 (2020).

7. Weigelt, P., König, C. & Kreft, H. GIFT – A Global Inventory of Floras and Traits for macroecology and biogeography. J. Biogeogr. 47, 16–43 (2020).

8. Enquist, B. J., Condit, R., Peet, R. K., Schildhauer, M. & Thiers, B. M. Cyberinfrastructure for an integrated botanical information network to investigate the ecological impacts of global climate change on plant biodiversity. https://peerj.com/preprints/2615 (2016) doi:10.7287/peerj.preprints.2615v2.

9. Wang, H. et al. The China plant trait database version 2. Sci. Data 9, 769 (2022).

10. Tavşanoğlu, Ç. & Pausas, J. G. A functional trait database for Mediterranean Basin plants. Sci. Data 5, 180135 (2018).

11. Falster, D. et al. AusTraits, a curated plant trait database for the Australian flora. Sci. Data 8, 254 (2021).

12. Báez, S. et al. FunAndes – A functional trait database of Andean plants. Sci. Data 9, 511 (2022).

13. Kissling, W. D. et al. PalmTraits 1.0, a species-level functional trait database of palms worldwide. Sci. Data 6, 178 (2019).

14. Klimešová, J. et al. Handbook of standardized protocols for collecting plant modularity traits. Perspect. Plant Ecol. Evol. Syst. 40, 125485 (2019).

15. WFO. The World Flora Online (WFO). http://www.worldfloraonline.org/ (2023).

16. Díaz, S. et al. The global spectrum of plant form and function. Nature 529, 167–171 (2016).

17. Garnier, E. et al. Towards a thesaurus of plant characteristics: an ecological contribution. J. Ecol. 105, 298–309 (2017).

18. Raunkiaer, C., Egerton, F. N. & Fausboll. The life forms of plants and statistical plant geography. (Clarendon Press, 1934).

19. Beentje, H. The Kew Plant Glossary: An Illustrated Dictionary of Plant Terms. (Kew Publishing, 2016).

20. Simpson, M. G. Plant Systematics. (Elsevier, 2019).

21. Cornelissen, J. H. C. et al. A handbook of protocols for standardised and easy measurement of plant functional traits worldwide. Aust. J. Bot. 51, 335 (2003).

22. Pérez-Harguindeguy, N. et al. New handbook for standardised measurement of plant functional traits worldwide. Aust. J. Bot. 61, 167–234 (2013).

23. Saatkamp, A. et al. A research agenda for seed-trait functional ecology. New Phytol. 221, 1764–1775 (2019).

24. Wigley, B. J. et al. A handbook for the standardised sampling of plant functional traits in disturbance-prone ecosystems, with a focus on open ecosystems. Aust. J. Bot. 68, 473– 531 (2020).

25. Freschet, G. T. et al. A starting guide to root ecology: strengthening ecological concepts and standardising root classification, sampling, processing and trait measurements. New Phytol. 232, 973–1122 (2021).

26. Ely, K. S. et al. A reporting format for leaf-level gas exchange data and metadata. Ecol. Inform. 61, 101232 (2021).

27. Kleyer, M. et al. The LEDA Traitbase: a database of life-history traits of the Northwest European flora. J. Ecol. 96, 1266–1274 (2008).

28. Sauquet, H. et al. The ancestral flower of angiosperms and its early diversification. Nat. Commun. 8, 16047 (2017).

29. Lenters, T. P. et al. Integration and harmonization of trait data from plant individuals across heterogeneous sources. Ecol. Inform. 62, 101206 (2021).

30. Arnaud, E. et al. Towards a reference Plant Trait Ontology for modeling knowledge of plant traits and phenotypes. in Proceedings of the International Conference on Knowledge Engineering and Ontology Development (KEOD-2012), 220--225 (2012). doi:10.5220/0004138302200225.

31. Schentz, H., Peterseil, J. & Bertrand, N. EnvThes – interlinked thesaurus for long term ecological research, monitoring, and experiments. in Proceedings of the 27th Conference on Environmental Informatics - Informatics for Environmental Protection, Sustainable Development and Risk Management. (Page, B., Fleischer, A. G., Göbel, J. & Wohlgemuth, V. (Hrsg.), 2013).

32. Schneider, F. D. et al. Towards an ecological trait-data standard. Methods Ecol. Evol. 10, 2006–2019 (2019).

33. Kattge, J. et al. A generic structure for plant trait databases. Methods Ecol. Evol. 2, 202– 213 (2011).

34. Madin, J. S. et al. The Coral Trait Database, a curated database of trait information for coral species from the global oceans. Sci. Data 3, 160017 (2016).

35. Wilkinson, M. D. et al. The FAIR Guiding Principles for scientific data management and stewardship. Sci. Data 3, 160018 (2016).

36. Riley, J. Understanding Metadata: What is Metadata, and What is it For?: A Primer. https://www.niso.org/publications/understanding-metadata-2017 (2017).

37. Miles, A. & Bechhofer, S. SKOS Simple Knowledge Organization System Reference. https://www.w3.org/TR/skos-reference/ (2009).

38. Brickley, D. & Guha, R. V. RDF Schema 1.1. W3C. 1.1. RDF Working Group https://www.w3.org/TR/rdf-schema/ (2014).

39. DCMI Metadata Terms. https://www.dublincore.org/specifications/dublin-core/dcmi-terms/2010-10-11/ (2010).

40. Schadow, G., McDonald, C. J., Suico, J. G., Föhring, U. & Tolxdorff, T. Units of Measure in Clinical Information Systems. J. Am. Med. Inform. Assoc. 6, 151–162 (1999).

41. Dumontier, M. et al. The Semanticscience Integrated Ontology (SIO) for biomedical research and knowledge discovery. J. Biomed. Semant. 5, 14 (2014).

42. Madin, J. S. et al. An ontology for describing and synthesizing ecological observation data. Ecol. Inform. 2, 279–296 (2007).

43. DataCite Metadata Schema 4.4. DataCite Schema https://schema.datacite.org/meta/kernel-4.4/ (2021).

44. Magagna, B. et al. The I-ADOPT Interoperability Framework: a proposal for FAIRer observable property descriptions. https://w3id.org/iadopt/ont/0.9.1 (2021) doi:10.5194/egusphere-egu21-13155.

45. Boettiger, C. rdflib: A high level wrapper around the redland package for common rdf applications (Version 0.1.0). (2023).

46. Spjut, R. W. A Systematic Treatment of Fruit Types. The World Botanical Associates http://www.worldbotanical.com/fruit_types.htm (2015).

47. Díaz, S. et al. The global spectrum of plant form and function: enhanced species-level trait dataset. Sci. Data 9, 755 (2022).

48. Mayfield, E. Illustrated Plant Glossary. (CSIRO Publishing, 2021).

49. Harris, J. G. & Harris, M. W. Plant Identification Terminology: An Illustrated Glossary. (Spring Lake Pub., 2001).

50. Wäldchen, J., Wittich, H. C., Rzanny, M., Fritz, A. & Mäder, P. Towards more effective identification keys: A study of people identifying plant species characters. People Nat. 4, 1603–1615 (2022).

51. anonymous. Systematics Association Committee for Descriptive Biological Terminology, IIa. Terminology of Simple Symmetrical Plane Shapes (Chart 1a), Addendum. Taxon 11, 245–247 (1962).

